# Unsupervised segmentation of 3D microvascular photoacoustic images using deep generative learning

**DOI:** 10.1101/2023.04.30.538453

**Authors:** Paul W. Sweeney, Lina Hacker, Thierry L. Lefebvre, Emma L. Brown, Janek Gröhl, Sarah E. Bohndiek

## Abstract

Mesoscopic photoacoustic imaging (PAI) enables label-free visualisation of vascular networks in tissue at high contrast and resolution. The segmentation of vascular networks from 3D PAI data and interpretation of their meaning in the context of physiological and pathological processes is a crucial but time consuming and error-prone task. Deep learning holds potential to solve these problems, but current supervised analysis frameworks require human-annotated ground-truth labels. Here, we overcome the need for ground-truth labels by introducing an unsupervised image-to-image translation deep learning model called the *vessel segmentation generative adversarial network* (VAN-GAN). VAN-GAN integrates synthetic blood vessel networks that closely resemble real-life anatomy into its training process and learns to replicate the underlying physics of the PAI system in order to learn how to segment vasculature from 3D biomedical images. With a variety of *in silico, in vitro* and *in vivo* data, including patient-derived breast cancer xenograft models, we show that VAN-GAN facilitates accurate and unbiased segmentation of 3D vascular networks from PAI data volumes. By leveraging synthetic data to reduce the reliance on manual labelling, VAN-GAN lowers the barrier to entry for high-quality blood vessel segmentation to benefit users in the life sciences applying PAI to studies of vascular structure and function.

## Introduction

Mesoscopic photoacoustic imaging (PAI) provides high-resolution, contrast-rich images of vascular structures *in vivo* based on the absorption of light by haemoglobin. These images reveal subtle changes in vascular architecture in tissues, which have demonstrated importance in diagnosis and staging of a range of diseases, from diabetes to oncology ^1,2^. To quantify disease status, segmenting microvascular networks from 3D image volumes is vital, however, the development of segmentation frameworks has not kept pace with the rapid advancements in imaging technology. While deep learning offers a promising solution ^3^, its effective application in this field is fraught with complexities. Current supervised segmentation frameworks ^4–6^ typically require matched pairs of image volumes and human-annotated ground-truth feature masks. Manually generating vascular labels is time-consuming so is typically limited to a small number of image pairs. Furthermore, human annotation is error-prone, particularly in complex pathological tissues like tumours^7,8^ or in the presence of imaging artefacts, where interpretation of what constitutes a blood vessel can vary between experts. Supervised blood vessel segmentation models have been demonstrated based on annotated datasets in 2D retinal fluorescein angiograms ^9,10^ or 3D multiphoton images of brain vasculature ^11–13^. Efforts have also been made to develop semi-supervised methods, which reduce the paired dataset size required ^14–18^ or minimise ground-truth ambiguity ^18^. Given the challenges associated with manual labelling, unsupervised learning has naturally gained special attention for image segmentation tasks ^19^. In particular, cycle-consistent generative adversarial networks (CycleGAN) can be used to transform images from a imaging (source) domain to a segmentation (target) domain. By assuming an underlying relationship between each domain, CycleGAN performs style transfer between datasets of unpaired images by allowing two neural network translators to be trained in a constrained unsupervised and competitive manner. Modified formulations of CycleGAN have accurately segmented biomedical images ^20–24^, including cell nuclei from 3D fluorescence microscopy ^25^ and blood vessels in 2D X-ray angiograms ^26^.

To date, the application of deep learning methods in PAI has primarily focused on image reconstruction ^27–29^ or bridging the domain gap between simulations and experiments ^30,31^. Supervised methods have not only explored the task of blood vessel segmentation from 2D photoacoustic images ^32–36^ but have also shown promise in addressing the complexities involved in segmenting 3D images, as recent work has demonstrated the potential of combining synthetic data and manual annotations for effective supervised segmentation of 3D photoacoustic images ^37^. However, the transition to unsupervised methods in 3D PAI has been challenging due to artefacts arising from low signal-to-noise ratio (SNR) ^38^ or excitation and detection geometries ^39^, which severely limit their effectiveness. Whilst innovative unsupervised approaches have been applied in other 3D imaging modalities ^40,41^, there remains a need for tailored unsupervised models which overcome specific challenges in 3D PAI.

In this study, we introduce VAN-GAN (*<u>V</u>essel Segment<u>a</u>tio<u>n</u> <u>G</u>enerative <u>A</u>dversarial <u>N</u>etwork*), an innovative unsupervised deep learning model tailored to 3D vascular network segmentation in mesoscopic PAI. Distinctly surpassing traditional supervised approaches, VAN-GAN eliminates the need for human-annotated labels, employing image-to-image translation to adeptly transform PAI volumes into precise 3D vascular segmentation maps. Our model leverages a synthetic dataset enriched with a variety of vascular morphologies, further enhanced by advanced 3D deep residual U-Net generators and cycle-consistency loss functions. A significant breakthrough of VAN-GAN is its ability to segment vascular networks from tissue types beyond those included in its training dataset, showcasing adaptability and precision even in low signal-to-noise ratio environments typical of PAI. Demonstrating unparalleled performance on diverse *in silico* and *in vivo* PAI datasets, VAN-GAN not only outperforms conventional methods but also exhibits remarkable robustness against user bias and common PAI artefacts.

## Results

### Advancing 3D vessel segmentation in photoacoustic imaging using VAN-GAN

VAN-GAN is engineered to learn mappings between the *imaging domain* and *segmentation domain*, thereby training a segmenter that is adept for real-world PAI applications (Fig. 1A). The imaging domain comprises PAI volumes, either generated through physics-driven simulations or acquired experimentally. The segmentation domain consists of computer-generated branching blood vessel structures in 3D, stochastically created based on mathematical formalisms (see Methods). Throughout our study, the dataset for the segmentation domain remains consistent, and during the training of VAN-GAN, the datasets for imaging and segmentation domains are treated as unpaired.

**Figure 1.**
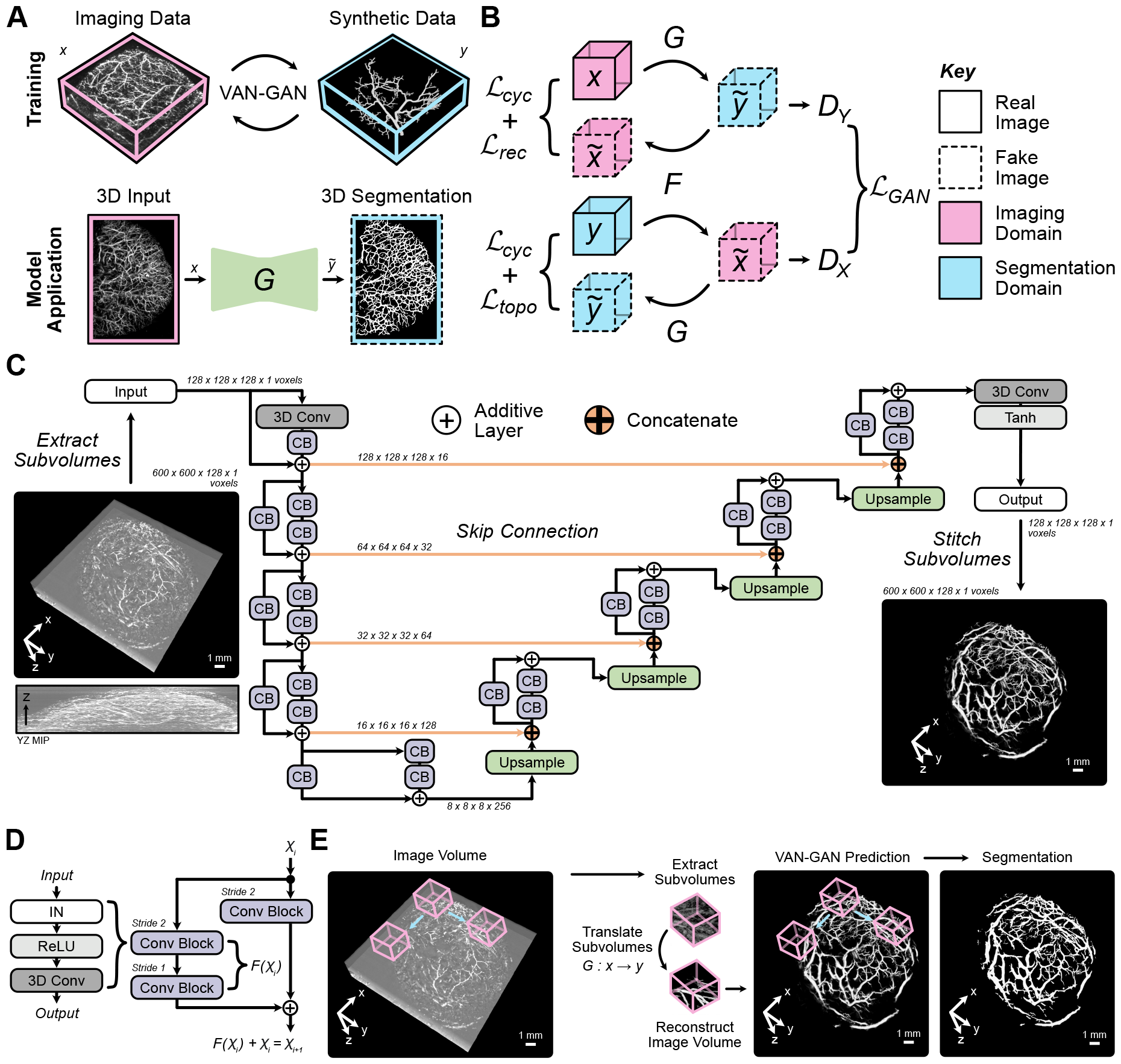
Vessel segmentation generative adversarial network (VAN-GAN) architecture for unsupervised segmentation of blood vessels in photoacoustic imaging (PAI) volumes. (A) The training and application process of VAN-GAN utilises two unpaired datasets (real PAI volumes, *x* and synthetic blood vessels, *y*) to train the segmentor, *G*, for real-world use. (B) VAN-GAN adapts the cycleGAN model and learns mappings between imaging (*x*) and segmentation (*y*) domains 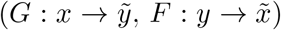 using additional reconstruction, *ℒ* _*rec*_, and topological, *ℒ*_*topo*_, cycle-consistency constraints. (C) A 3D deep residual U-Net generator architectures are employed. An example, PAI subvolume input (left) and segmentation output (right) is shown. (D) Standard residual units are employed for each generator. (E) A sliding window approach is used to form a probability map that is binarised to create the final segmentation mask. CB = convolutional block; MIP = maximal intensity projection; IN = instance normalisation.

VAN-GAN builds upon the CycleGAN model ^42^ with several key enhancements (see Methods and Supplementary Methods for full details). The generator architecture integrates U-Net ^4^ with deep residual learning ^43^ (Supplementary Table 1), and random Gaussian noise is introduced to the discriminator inputs and layers for improved training stability and regularisation (Supplementary Table 2). VAN-GAN is adapted to process 3D image volumes using 3D convolutions. It incorporates cycle-consistency loss functions to improve performance: a structural similarity reconstruction loss, which is applied to both real and reconstructed images in the imaging domain; and a spatial and topological constraint, applied to the real and reconstructed images in the segmentation domain to preserve tubular structures ^44^ (Fig. 1B).

The generators consists of three parts: encoder, bridge, and decoder ^45^ (Fig. 1C). The encoder and decoder feature four layers with residual units (Fig. 1D) connected by skip connections. The convolutional block sequence includes instance normalisation ^46^, ReLU activation ^47^, and 3D convolution, with reflection padding to minimise boundary artefacts ^42,48^. VAN-GAN is trained on subvolumes of images, with predictions combined using a sliding window for full volume segmentation (Fig. 1E, see Methods).

### VAN-GAN rivals supervised learning in segmenting photoacoustic images

We first tested VAN-GAN using physics-driven simulations to virtually emulate PAI, giving the stochastic 3D branching vessel structures from VAN-GAN’s segmentation domain as simulation inputs (see Methods, Supplementary Methods, Supplementary Fig. 1 and 2). In essence, this created a paired *in silico* dataset of PAI volumes and their segmentations, which enables direct validation of VAN-GAN’s segmentation performance (Fig. 2A).

**Figure 2.**
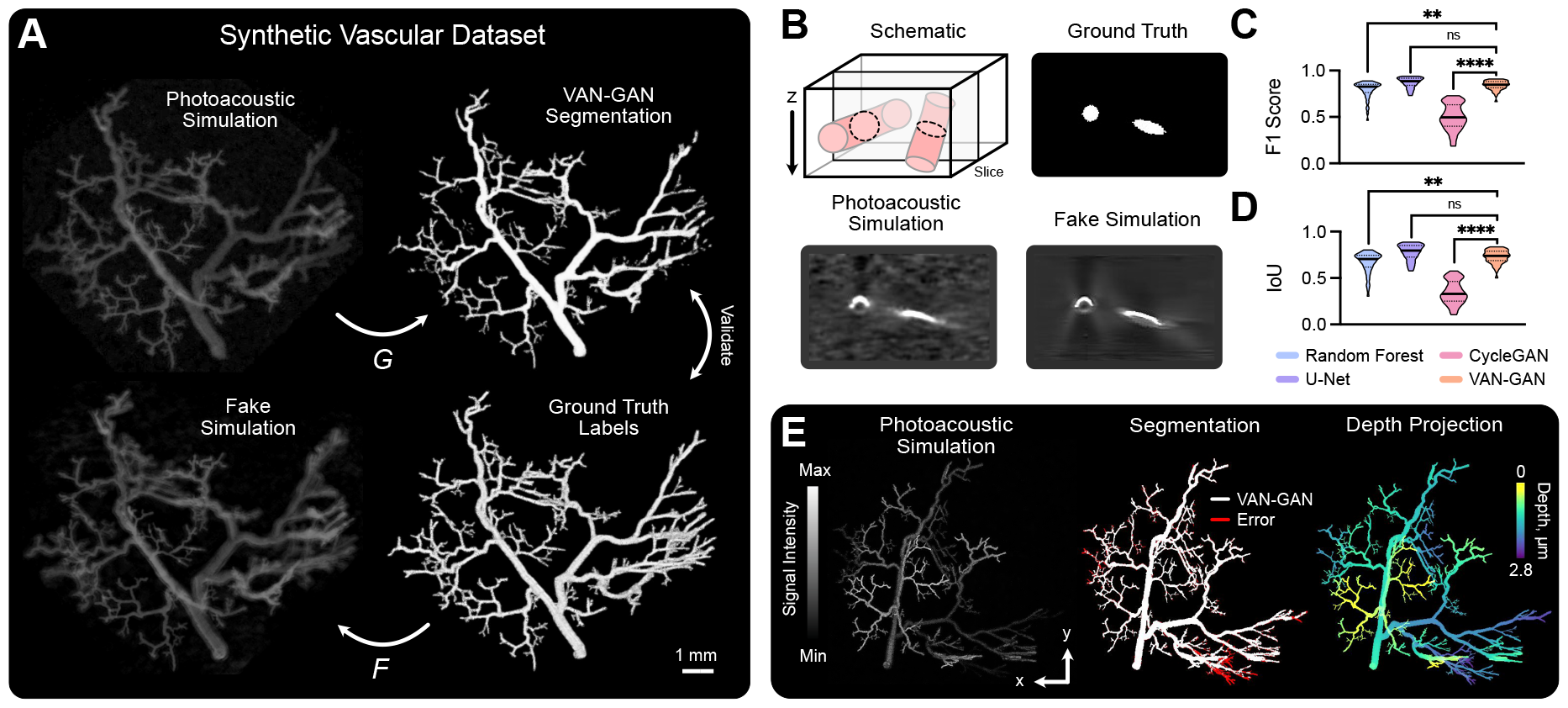
VAN-GAN performance on synthetic photoacoustic data. (A) Illustration of the paired dataset, consisting of synthetic vessel structures that form ground-truth labels and corresponding physics-driven PAI simulations. (B) Fake simulation images generated by generator *F* reproduce artefacts (present in our simulations) that are inherent to mesoscopic PAI and influence vessel shape and continuity. The example illustrates an artefact arising from limited angular coverage of the illumination fibres, which means the circular vessel cross-section appears as an arc in PAI. Segmentation metric distributions compare performance of a random forest pixel classifier, U-Net, CycleGAN and VAN-GAN:(C) F1 Score; and (D) Intersection over Union (IoU). (E) An example of (left) a photoacoustic simulation, (middle) the corresponding VAN-GAN segmentation labels and error with respect to the ground-truth, and (right) a colour-coded depth projection of the network. Data in (C-D) are represented by truncated violin plots with interquartile range (dotted) and median (solid) shown. Statistical significance indicated by * (*P <* 0.05), ** (*P <* 0.01),*** (*P <* 0.001) and **** (*P <* 0.0001). *ns* = no significance.

During training, VAN-GAN perceived images from both the imaging and segmentation domains as unpaired and thus processed them in an unsupervised manner. Then during testing, the paired dataset enabled us to perform an ablation study to quantitatively evaluate the impact of key VAN-GAN components on the segmentation accuracy via their systematic removal from the model. The ablation study indicated that VAN-GAN was able to generate fake PAI volumes that included typical photoacoustic artefacts, such as those arising from limited illumination and detection views, as well as shadowing from overlying absorbers ^39^ (Fig. 2B). We found that omission of discriminator noise and reconstruction loss in the reduced models impeded learning of these image features, whereas the addition of the topological loss provides a greater constraint to improve segmentation of vascular structures (Table 1). The full VAN-GAN model achieved the highest F1 Score (0.842) and IoU (0.730) for the paired dataset.

**Table 1:**
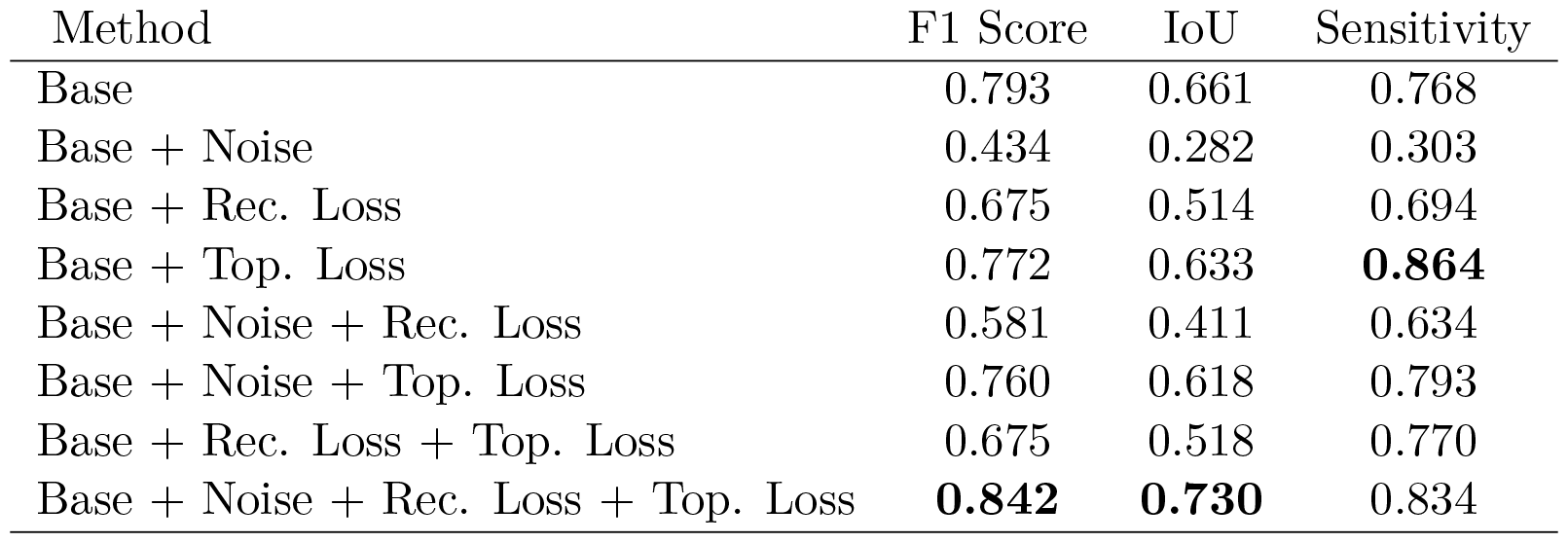
VAN-GAN ablation study. The following components were systematically removed to evaluate the performance of our model in their absence: 1) random Gaussian noise added to discriminator inputs and convolution layers (*Noise*); 2) a perceptual constraint on cycle reconstructed photoacoustic images (*Rec*.); and 3) a spatial and topological constrained on cycle reconstructed segmentation images (*Topo*.). In the *Base* setup, all these components are ablated. The inclusion of all three components, *Base + Noise + Rec. Loss + Top. Loss*, gives the full VAN-GAN model. Note, specificity excluded due to ∼ 0.999 values for all methods.

VAN-GAN was then benchmarked using the paired synthetic data against other deep-learning models selected for their relevance and proven effectiveness in image segmentation within the scope of PAI. Supervised methods included a random forest pixel classifier (RF, embeded within open-source package ilastik ^49^) and a 3D U-Net ^5^ (Supplementary Table 3). A comparison with CycleGAN ^42^ was also made (Supplementary Table 4), given its architectural similarities with VAN-GAN. CycleGAN and VAN-GAN treated the dataset as unpaired during training, whereas full feature labels were supplied to the RF and U-Net (see Supplementary Methods). The dataset consisted of 449 paired images with 10% assigned for testing and the remaining 90% split between training (80%) and validation (20%).

The results show good performance by VAN-GAN with, for example, mean F1 Score / IoU of 0.842 / 0.778, compared to the RF (0.792 / 0.665), U-Net (0.873 / 0.778) and CycleGAN (0.502 / 0.346) (Table 2). Significant differences between metrics were found when comparing VAN-GAN to RF and CycleGAN (Fig. 2C,D) and due to its poor performance, CycleGAN was excluded from all further analyses. Comparison to U-Net found there was no significant difference in performance in these fully labeled synthetic datasets, however, in an *in vivo* context, ground-truth annotations are partial at best and we found the U-Net trained on synthetic data lacked generalisability to *in vivo* data (Supplementary Fig. 3), hence the U-Net model was also excluded from all further analyses. RF methods are typically able to maintain performance with only partially labelled data, so RF was used as a comparator in subsequent analyses in the *in vivo* setting.

**Table 2:** Comparison of segmentation performance using PAI simulations. F1 Score, intersection over union (IoU), sensitivity and specificity were calculated on the testing dataset. RF = random forest pixel classifier and the highest metric score per ground-truth is shown in bold. Statistical significance between VAN-GAN is indicated by * (*P <* 0.05),** (*P <* 0.01), *** (*P <* 0.001) and **** (*P <* 0.0001). If blank, no significance was found compared to VAN-GAN.

| Dataset | Method | F1 Score | IoU | Sensitivity | Specificity |
| --- | --- | --- | --- | --- | --- |
| **Simulated** | RF | 0.792** | 0.665** | 0.748** | 0.999 |
|  | U-Net | **0.873** | **0.778** | **0.877** | **0.999** |
|  | CycleGAN | 0.502**** | 0.346**** | 0.558**** | 0.997**** |
|  | VAN-GAN | 0.842 | 0.730 | 0.834 | 0.999 |

Qualitative examination of VAN-GAN segmentation predictions showed errors were largely confined to the smallest vessels, which exhibited relatively low SNR as they were furthest from the simulated light-source in the tissue (Fig. 2E).

### VAN-GAN avoids user bias arising from human annotations

Given the need for human annotations in supervised training, we sought to compare the impact of expert variability on this process. Firstly, two experts independently labelled blood vessels on 2D maximum intensity projections (MIPs) generated from 3D *in vivo* PAI data from mouse ears (n=1) and skin (n=4) (Fig. 3A). MIPs were specifically employed to mitigate prevalent surface illumination artefacts that typically lead to only the upper part of absorbing structures being excited and detected, which has been shown to result in inaccurate 3D segmentations by human annotators ^38^.

**Figure 3.**
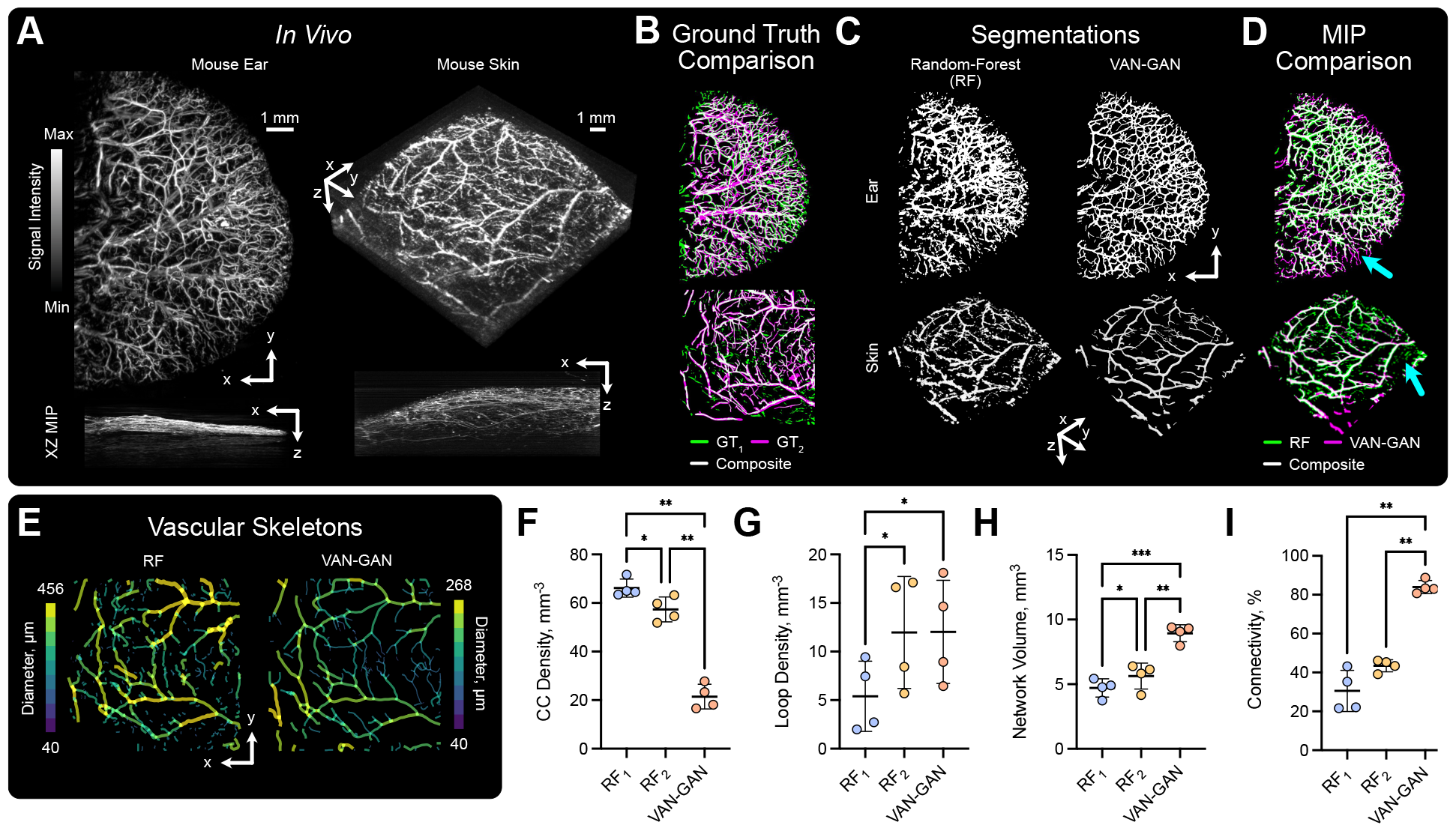
Unsupervised vascular segmentation overcomes user variation in human annotations. (A) Mouse ear and skin imaged *in vivo* using PAI. (B) ground-truths for XY maximal intensity projections (MIPs) were generated by two experts (indicated by subscripts *1* and *2*). Visualisations indicate variability between each expert for the ear (top) and skin (bottom). (C) 3D segmentation masks predicted using RF (left) and VAN-GAN (right) for datasets shown in (A,B). (D) Comparison of MIPs generated from the 3D segmentation labels shown in (C) with arrows indicating segmented noise and vessel discontinuities using RF. (E) 2D projections of 3D vascular skeletons created from the two segmentation methods applied to mouse skin. Variability between individual users RF classifiers and VAN-GAN on the skeletonised mouse skin is indicated for a range of statistical and vascular descriptors: (F) connected component (CC) density (the number of subnetworks normalised against tissue volume, mm ^− 3^); (G) loop density (the number of vessel network loops normalised against tissue volume, mm ^− 3^); (H) network volume (mm^3^); and (I) connectivity (the volume of the largest subnetwork normalised against total network volume, %);. Mean and standard deviation of data are shown in (F-I). Statistical significance indicated by * (*P <* 0.05), ** (*P <* 0.01), *** (*P <* 0.001) and **** (*P <* 0.0001). MIP = maximal intensity projection.

Comparing the two expert annotations, substantial differences were observed between their segmentations both qualitatively (Fig. 3B) and quantitatively (F1 Score: 0.667/0.569, IoU: 0.501/0.397, Specificity: 0.916/0.892 and Sensitivity: 0.754/0.659 for ear / skin), highlighting the challenges in ground-truth label creation in real imaging data. These expert annotators then independently trained separate 3D RF models for vessel segmentation in mouse ear and skin datasets. Comparing these models showed no bias toward respective expert labels; each RF model performed poorly against the 2D annotations (for example, F1 Score: 0.668 / 0.613 for ear / skin for the RF model and ground-truths created by expert 1 - Supplementary Table 5). VAN-GAN’s accuracy was also low overall (F1 Score: 0.649 / 0.529 for ear / skin compared against ground-truths labelled by expert 1), however, it was more robust to segmentation of background noise as closer scrutiny revealed limitations in RF, such as artefact segmentation and discontinuities in segmented vessels (Fig. 3C, D).

To analyse more comprehensively the 3D morphology of vascular networks segmented by the RF and VAN-GAN models, we skeletonised the 3D vessel masks (Fig. 3E) and calculated a set of vascular descriptors based on topological data analysis (see Methods). Focusing on the mouse skin, we found that the number of vascular subnetworks (or connected components) and network loops between RF_1_ and RF_2_ were significantly different (Fig. 3F,G). These analyses highlight a critical flaw in using supervised segmentation models - their performance is inherently limited by the quality of training data. Independent training by different expert users can lead to systematically varied segmentation outcomes (Fig. 3H), which could bias study interpretation. By comparison, VAN-GAN predicts more larger interconnected vascular networks, with the largest subnetwork on average forming 84.0% of the total subnetwork volume (Fig. 3H,I). These results indicate that VAN-GAN segments more well-connected networks compared to RF, which aligns better with the expectations of the biological network in healthy skin.

### VAN-GAN is more robust to artefacts compared to human annotations

Several imaging artefacts impede segmentation performance in PAI. Illumination artefacts, resulting from limited light coverage, cause objects to appear flattened in images (Fig. 4A), leading human users to annotate only the brighter top surfaces of blood vessels. In our paired synthetic dataset, we addressed this by training the supervised RF with full ground-truth labels, enabling it to learn the complete range of signal intensities expected from the reference structures. Training an RF segmenter with these paired synthetic data resulted in more accurate, axisymmetric vessel segmentation than that achieved by an RF model specialised for experimental *in vivo* datasets and trained by a human annotator (Fig. 4B,C).

**Figure 4.**
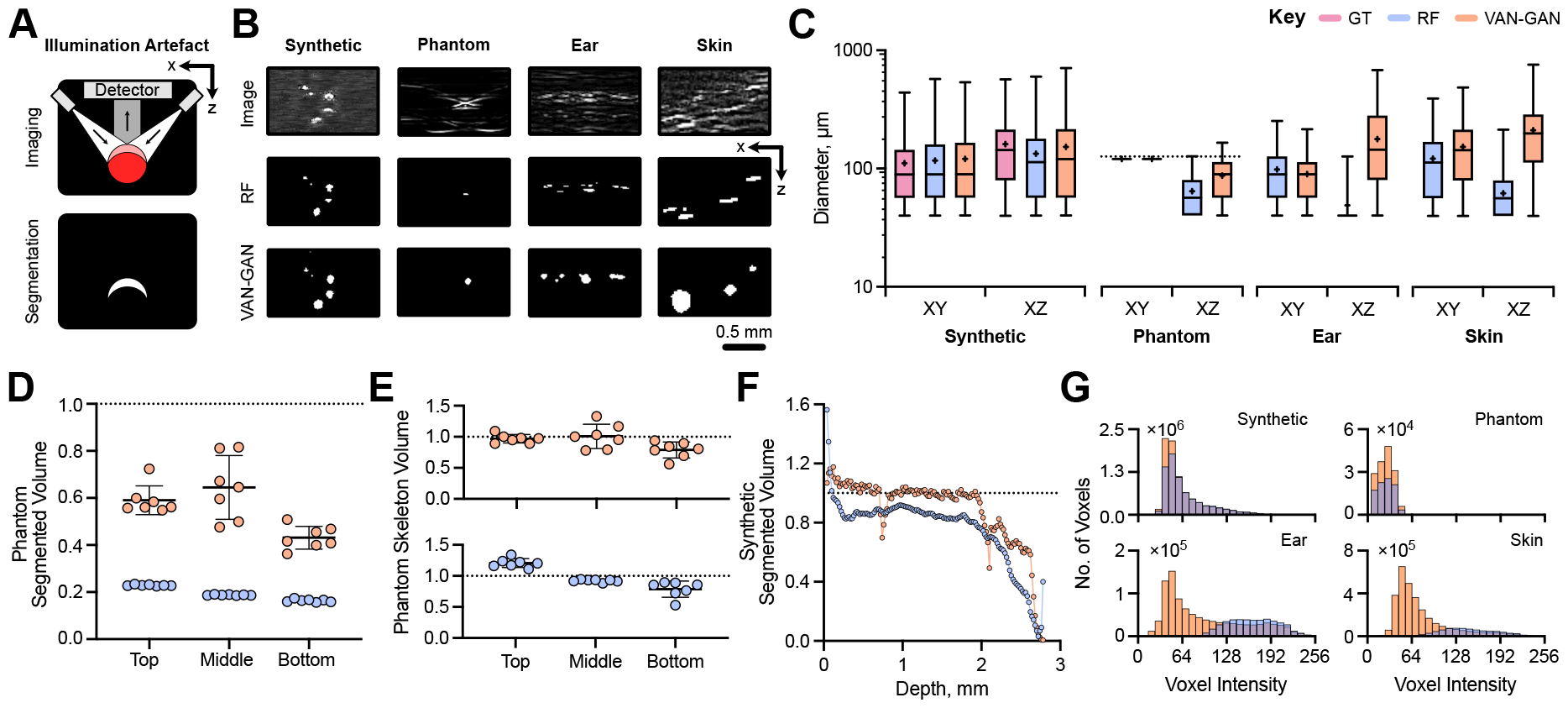
VAN-GAN overcomes image artefacts in photoacoustic mesoscopy. (A) The illumination artefact in photoacoustic mesoscopy results in the incorrect segmentation of the top surface of blood vessels due to lower signals from underlying regions. (B) Example image subsections highlighting the illumination artefact in each imaging domain dataset and corresponding random forest pixel classifier (RF) and VAN-GAN segmentations. RF models for phantom, ear and skin segmentation were trained by an expert whereas the RF model for the synthetic data was trained using known computer-generated groundtruths. (C) Diameters in the XY and XZ directions calculated for each dataset. For the string phantom, the known string volume is indicated by the dotted line. Computed string phantom volumes from (D) 3D segmented images and (E) skeletonised strings normalised against the ground-truth value. (F) Normalised segmented volume for our synthetic dataset with respect to tissue depth. (G) Histograms of signal intensities for each voxel segmented by each method and for each dataset: synthetic vasculatures, *in situ* string phantoms and *in vivo* mouse ear and skin datasets. Data in (C) is given by box and whisker plots which display the 25 ^th^, 50 ^th^ and 75 ^th^ quartiles, and the interquartile range. Mean and standard deviation of data are shown in (D) and (E).

In contrast, VAN-GAN effectively accounts for the entire signal intensity range across vessel walls and lumens in both *in silico* and *in vivo* datasets. When comparing diameters in the XZ-plane, VAN-GAN’s predictions were closer to ground-truth, outperforming the RF model (Fig. 4B,C). Accuracy was further validated using photoacoustic string phantoms, which consist of three non-overlapping strings of known size located at three separate depths parallel to the imaged surface ^39^. Although VAN-GAN was not specifically trained on string phantom images, it more accurately predicted string diameters in the XZ-plane than the RF model, demonstrating its robustness and versatility to illumination artefacts.

Further artefacts arise in PAI due to depth-dependent SNR. VAN-GAN demonstrated a consistent ability to accurately predict string volumes at various depths (Fig. 4D). Both VAN-GAN and RF models showed improved network volume predictions after applying skeletonisation, which presumes axisymmetric vessels (Fig. 4E). On the synthetic dataset, VAN-GAN consistently outperformed RF, especially when depth exceeded 2 mm (Fig. 4F). A detailed analysis of segmented voxel signal intensities revealed a tendency for VAN-GAN to segment more low-intensity voxels across all datasets, a trend that became more pronounced for *in vivo* data (Fig. 4G). Compared to RF models with their limited receptive fields, VAN-GAN appears superior in handling complex spatially-varying background noise and can better learn intricate feature representations.

### VAN-GAN segments vascular topologies beyond the training dataset

PAI is vital for monitoring blood vessel evolution in tumours, which present unique segmentation challenges due to their heterogeneous nature ^50,51^. Unlike physiological tissues, tumour vascular architecture is chaotic with varying diameters, lengths and inter-connectivity across various spatial scales. These features are absent in VAN-GAN’s synthetic segmentation domain dataset since the method used to generate branching structures inherently leads to regular and predictable patterns.

To evaluate the ability of VAN-GAN to segment complex pathological vascular networks, datasets of 3D images of oestrogen receptor positive (ER+) and negative (ER-) breast cancer tumours derived from both patient-derived xenograft models ^38,52^ and cell lines (MCF7 and MDA-MB-231, respectively) were used. Segmentations made by VAN-GAN allowed hypothesised structural differences in vasculature between the ER+ and ER-subtypes to be identified in both tumour types (Fig. 5A and Supplementary Notes). In the PDX tumours, the ER-tumours exhibited significantly higher vessel surface area density with respect to tumour volume (*P <* 0.05, Fig. 5B). The *ex vivo* immunohisto-chemistry (IHC) analysis of CD31 staining, an endothelial cell marker, cross-validated this finding, as the ER-tumours showed significantly greater CD31 positivity (*P <* 0.05, Fig. 5C). Additionally, VAN-GAN indicated ER-tumours displayed a higher density of vascular looping structures (*P <* 0.01, Fig. 5D) and reduced vessel lengths (*P <* 0.01), which could indicate a more immature vascular network compared to ER+. IHC staining of *α*-smooth muscle actin (*α*SMA), a pericyte and smooth muscle marker, colocalised with CD31, supported this finding as ER-tumours showed significantly lower positivity (*P <* 0.01, Fig. 5E).

**Figure 5.**
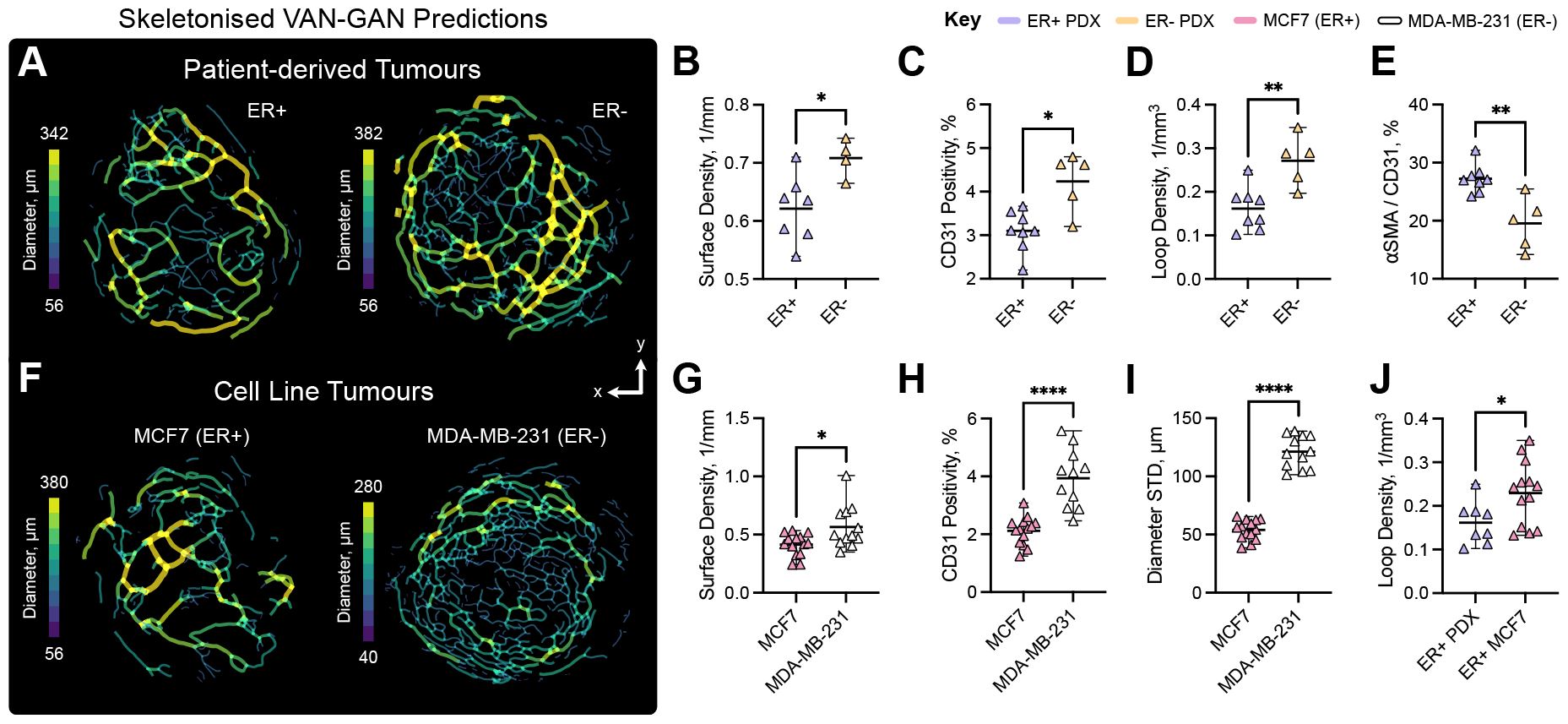
VAN-GAN enables quantification of pathological vascular architecture. (A) Vascular skeletons for oestrogen receptor negative (ER-, left) and positive (ER+, right) breast cancer patient-derived xenograft tumours. A comparison of metrics for ER- (top) and ER+ (bottom): (B) vessel surface density (1/mm), (C) CD31 positivity (staining area with respect to tumour area, %), (D) vessel loop density (1/ mm^3^) and (E) *α*-smooth muscle actin (*α*SMA) colocalisation with CD31 staining (%). (F) Vascular skeletons of MCF7 (left) and MDA-MB-231 (right) breast cancer tumours. VAN-GAN metrics for MCF7 and MDA-MB-231 tumours: (G) vessel surface density (1/mm), (H) CD31 positivity and (I) standard deviation (STD) of vessel diameters (*μ*m). (J) A comparison of vessel loop density (1/ mm^3^) in ER+ models. Mean and standard deviation of data are shown in (C-F) and (H-K). Statistical significance indicated by * (*P <* 0.05), ** (*P <* 0.01), *** (*P <* 0.001) and **** (*P <* 0.0001).

VAN-GAN also showed distinct features between the cell-line derived MCF7 (ER+) and MDA-MB-231 (ER-) tumours (Fig. 5F) that were cross-validated by IHC, where the percentage of vessel wall surface was quantified on CD31 stained sections with respect to intra- and extra-vascular area. Here, when quantifying blood vessel network surface area with respect to the tumour volume from our 3D VAN-GAN segmentations, similar trends were observed (*P <* 0.05, Fig. 5G), with ER-MDA-MB-231 tumours displaying significantly elevated levels of CD31 (*P* = 0.0001, Fig. 5H). *α*SMA staining also indicated lower positivity for ER-MDA-MB-231 tumours (*P <* 0.01) but no significance in vessel loops or lengths between groups was found between each oestrogen receptor group. Vessels in the ER-MDA-MB-231 tumours exhibited greater heterogeneity (vessel diameters standard deviation, *P <* 0.0001, Fig. 5I), in contrast to the pattern observed in PDX tumours where greater heterogeneity was noted in ER+ tumours (*P <* 0.05). Observed differences between PDX and cell-line derived tumours were underscored by a significant difference in looping structure between ER+ models (*P <* 0.05, Fig. 5J), highlighting divergence in vascular maturity. The ER+ PDX tumours were likely at a more advanced stage of vascular development compared to ER+ MCF7 tumours.

The close relationship of the topological descriptors extracted from VAN-GAN segmentations to IHC analyses of the same tumours, together with common trends across different tumour models, indicate that VAN-GAN is able to accurate segment complex vasculatures that exceed the constraints imposed by the synthetic segmentation data used in its training.

## Discussion

Here, we introduced VAN-GAN, an innovative deep learning model that segments 3D vascular networks imaged using PAI mesoscopy. For mesoscopic PAI, human annotations are time consuming and laborious due to depth-dependent SNR and subtle imaging artefacts, which make it particularly challenging to label pathological tissues such as tumours. Independent ground-truth labelling by two expert users in our study showed substantial discrepancies in both their segmentations and the vascular topology parameters measured from subsequent skeletonisations, highlighting the potential for detrimental impact of user bias on quantification of blood vessel networks.

VAN-GAN adeptly navigates these difficulties in multiple ways. Firstly, VAN-GAN is a novel approach that builds on the foundation of CycleGAN by integrating 3D deep residual U-Net generators and bespoke cycle-consistency loss functions and discriminator noise, fully leveraging the power of unsupervised learning for PAI image segmentation. These additional elements were found through an ablation study to enhance the capability of VAN-GAN in segmenting intricate vascular structures from synthetic PAI volumes, leading to VAN-GAN surpassing traditional supervised methods and rivalling the gold standard U-Net.

Secondly, VAN-GAN is able to handle complex imaging artefacts arising from the geometry of the PAI system, demonstrating robustness in both synthetic and real-world datasets. VAN-GAN errors were generally confined to only the smallest vessels. Importantly, VAN-GAN provided realistic quantification of vessel lumens, which otherwise appear flattened in supervised segmentations due to illumination artefacts; VAN-GAN restored the segmenation that would be expected based on reference structures (in phantoms) and maintained lumen patency through to healthy ear and skin tissues. VAN-GAN also demonstrated greater robustness to depth-dependent SNR, segmenting at a greater depth than supervised methods and providing the most accurate quantifications with depth of the tested methods.

Finally, VAN-GAN provides biologically relevant segmentations for *in vivo* PAI data, showing more interconnected and larger vascular networks in the healthy ear and skin than other methods. VAN-GAN also extended directly to application in pathological tissues, such as patient-derived breast cancer, even though the complex chaotic architectures associated with these tissue types were absent from the synthetic dataset used for training. Topological data analysis of skeletonised vascular networks derived from VAN-GAN segmentations showed biological findings consistent with IHC analysis conducted on *ex vivo* sections. Taken together, these three main findings demonstrate strengths of VAN-GAN to significantly boost the precision and reliability of vascular segmentation in PAI, while emphasising the versatility of the approach.

VAN-GAN has demonstrated impressive capabilities, however, it is important to consider the limitations of its training. Reliance on synthetic data, though effective, may in future need further adaptation to encompass the full diversity of real-world vascular structures, particularly when considering application to human imaging data. Such adaptations could include developing more complex simulations to create more realistic vasculatures for simulation or integrating manual 3D labels from a diverse range of imaging techniques into the training data. Additionally, extending applicability to a wider range of tissue types and chromophores, beyond those included in its initial training, would be important. All of the animal studies undertaken here were in nude mice, which lack skin pigmentation, however, skin tone is a consideration that is gaining greater attention in the PAI community ^53^ and data from a range of skin tones would be needed to maximise applicability of VAN-GAN in future.

Moving forward, the potential applications of VAN-GAN extend beyond PAI. A key focus area for enhancement is the optimisation of training schemes and loss function weightings, which are crucial for ensuring the generalisability and efficacy of the model across diverse imaging contexts. Merging VAN-GAN with open-source bioimage platforms would also be important to democratise access to advanced segmentation tools, fostering wider adoption and application in the life sciences. Integration such as this not only aligns with the trend towards accessible, high-quality image analysis but also opens new avenues for research and clinical applications by providing consistent and unbiased results.

In conclusion, VAN-GAN sets a new precedent in the segmentation of 3D microvascular networks in mesoscopic PAI. By reducing the reliance on manual labelling and leveraging synthetic data, our approach promises to lower the barrier to entry for high-quality blood vessel segmentation, leading to more robust and consistent characterisation of vascular structures. VAN-GAN could thus not only deepen our understanding of tumour vascular architectures but also pave the way for the discovery of novel vascular-targeted therapeutics and improvement of diagnostic accuracy across clinical applications.

## Methods

The following details architecture and training methodology of VAN-GAN, in addition to describing image synthesis and preprocessing. VAN-GAN was implemented using Tensorflow ^54^ with Keras backend ^55^ and Tensorflow Addons, along with Tensorflow-MRI ^56^. The model was trained on either: 1) a Dell Precision 7920T with a Dual Intel Xeon Gold 5120 CPU with 128GB RAM and two NVIDIA Quadro GV100 32GB GPUs with NVLink; or 2) a custom built workstation with a Intel Xeon Gold 5220 CPU with 128GB RAM and four NVIDIA RTX A6000 48GB GPUs with NVLinks. The optimal model for each dataset was selected for application based on qualitative image evaluation of generated images from the test set and a comprehensive analysis of minima for each loss function (see Supplementary Methods and Supplementary Figs. 4-6).

### 3D Deep Residual U-Net Generators

VAN-GAN generators use a modified version of a deep residual U-Net architecture ^45^ (see Supplementary Table 2), which integrates residual units into a U-Net architecture to ease training and facilitate information propagation through the network without degradation. For the latter, it is important to use low-level details and retain high-resolution semantic information to improve segmentation performance ^4,45,57^. Both the generator input and output tensor shapes are 128 *×* 128 *×* 128 *×* 1 (*depth × width × height × channel*).

### 3D PatchGAN Discriminators

The discriminators use a five layer PatchGAN architecture ^58^ (see Supplementary Table 3). Each layer is composed of a 3D convolutional layer, instance normalisation ^46^, leaky ReLU ^59^ and spatial dropout ^60^ (a rate of 20% and excluded from first and final layers). Similarly to our generators, reflection padding was also used prior to convolution layers to reduce feature map artefacts ^48^. Further, for additional regularisation and to limit unstable behaviour of VAN-GAN during training, random Gaussian noise was added to real or fake inputs to the discriminator ^61,62^ and for every proceeding layer prior to convolution blocks ^62^ (Supplementary Table 4).

### Loss Functions

The goal is to learn the mapping functions *G* : *X* → *Y* and *F* : *Y* → *X* between the imaging, *X*, and segmentation, *Y*, domains. In the VAN-GAN model, each generator is designed to minimise its own respective cycle-consistency loss, rather than collectively minimising a total cycle-consistency loss as in CycleGAN ^42^. Consequently, for each domain transformation the corresponding generator is responsible for reducing the discrepancy between the original input and cycle reconstructed output. Separating total cycle consistency components enables more specialised optimisation of the generators by reducing the potential for conflicting domain transformation objectives, particularly given our imaging and segmentation domains are highly disparate.

We utilise *L*_1_-norm for forward cycle-consistency, i.e., *x* → *G*(*x*) → *F* (*G*(*x*)) ≈ *x*:

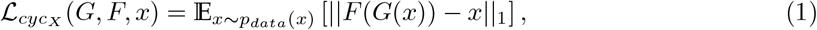

and binary cross-entropy for backward cycle-consistency, i.e., *y* → *F* (*y*) → *G*(*F* (*y*)) ≈ *y*:

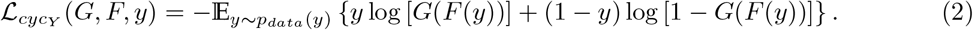

In addition to (1) and (2), two additional constraints on cycle-consistency are imposed: 1) a structural similarity index measure (SSIM) loss; and 2) a topology-preserving loss (centrelineDice or clDice ^44^) for backward cycle-consistency. SSIM is used for forward-consistency to ensure structural and perceptive features in biomedical images are retained when generating fake images. Similarly, to preserve the morphological characteristics of vascular networks when segmenting blood vessels, a constraint on backward-consistency was applied that seeks to minimise differences in network structure and topology.

SSIM is a perceptually motivated model composed of three comparison functions: luminance, contrast and structure. Typically SSIM is used to assess image quality, so as a loss function can be used for image restoration ^63^, such as denoising and super-resolution ^64,65^ and pose-guided person image generation ^66^. Here, SSIM adopts a sliding Gaussian window to calculate the SSIM index between two local patches centred around a pixel coordinate. For image patches *I*_*p*_ and *J*_*p*_, SSIM is defined as

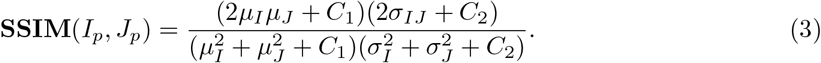

Here, *μ*_*I*_, *μ*_*J*_, 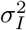 and 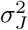 are the mean and variance of *I*_*p*_ and *J*_*p*_, respectively, *σ*_*IJ*_ is the covariance of *I*_*p*_ and *J*_*p*_, and *C*_1_ = (0.01 · *L*)^2^ and *C*_2_ = (0.03 · *L*)^2^, where *L* is the dynamic range of the pixel-values. To maximise the SSIM of biomedical images we form the reconstruction loss:

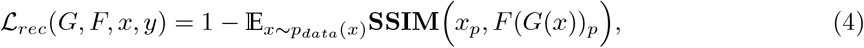

where the subscript *p* indicates image patches.

Cycle-consistency alone does not provide sufficient spatial constraint on the network topology of segmented images ^44^. Consequently, spatial and topological constraints on backward-consistency are added to act as additional regulatory loss function term. Here, the segmentation labels of synthetically-generated 3D vascular networks, *y*, are compared to *G*(*F* (*y*)) to ensure differences in topology are minimised. Minimisation is achieved using the connectivity-preserving metric **clDice**, which enforces topology preservation up to homotopy equivalence for binary segmentation^44^. Following Shit et al. ^44^, the loss function is a combination of soft-Dice loss and the differentiable form of **clDice**, *soft***clDice**:

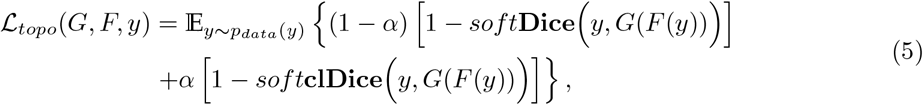

where the weighting, *α*, is set to 0.5.

The adversarial loss is expressed via least-squares adversarial loss ^67^ to mitigate problems with vanishing gradients. In the case of the mapping *G* : *X* → *Y*, this is given by

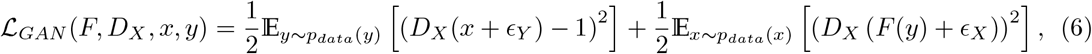

where *ϵ*_*X*_ and *ϵ*_*Y*_ are the randomly sampled Gaussian noise ^62^. Here the discriminator *D*_*X*_ aims to discern translated images, *F* (*y*), from real images *y* by minimising the objective function, whereas *F* aims to maximise it against its adversarial rival. Adversarial loss is similarly used for the mapping *G* : *X* → *Y*. Following Ihle et al. ^21^, we do not impose an identity mapping loss as this constrains the tint of an image ^42^ and so is not required for our greyscale and binary image volumes.

Thus, the objective function of VAN-GAN is given by

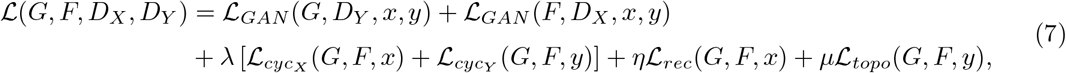

where the hyperparameters *λ, η* and *μ* control the relative importance between the objectives are set to 10, 5 and 5, respectively. We aim to solve:

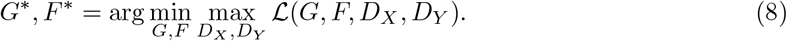

### Synthetic Vasculature

A language-theoretic model, called a Lindenmayer system (L-System) ^68^, was used to generate 3D branching vascular networks (termed here as *V-System*). L-Systems are ideally suited to our segmentation task as these models have been shown to create realistic, computer-generated 3D vascular branching structures ^69,70^ quickly and at scale ^38^ (O(10^2^) networks in O(1) minutes). To generate a synthetic branching network, we used a new stochastic grammar to create a string, which defines the complexity (in our case, the number of branching orders) of the vascular network, for example, branching order and angle, vessel diameter and tortuosity, and aneurysms or branching vessel shrinkage (see Supplementary Methods for mathematical descriptions). These strings are translated to graph form using a lexical and syntactic analyser and subsequently converted into volumetric binary segmentation masks ^38^, forming a synthetic dataset of 459 images for network training.

### Photoacoustic Simulations

To generate a paired image dataset, we performed photoacoustic simulations on the segmentation volume of each image in our synthetic vascular dataset. Each image pair consists of a physics-driven image volume and its corresponding known segmentation labels. Simulations followed the method of Brown et al. ^38^ who used SIMPA ^71^ (v0.1.1 with MCX v2020, 1.8) with the k-Wave MATLAB toolbox ^72^ (v1.3, MATLAB v2020b, MathWorks, Natick, MA, USA) to predict photoacoustic signals across synthetic vasculatures under the assumption that they are embedded in muscular tissue. In brief, vascular planar (XY) illumination was achieved on an isotropic resolution with optical forward modelling assuming an absorption spectrum of 50% oxygenated haemoglobin in blood vessels to mimic tumours ^73^. 3D acoustic forward modelling was then performed with the signal detected by a planar array of sensors positioned at the tissue surface, mimicking our PAI instrument (see below). The resulting photoacoustic wave-field was then reconstructed using a fast Fourier transform ^72^. While the PAI instrument raster-scans, we approximated this process with a planar illumination due to computational restrictions (reducing simulation time by a factor of 600^2^).

### Experimental Imaging

PAI was performed using a commercial system (Raster-scan optoacoustic mesoscopy RSOM Explorer P50, iThera Medical GmbH), as described previously ^38^. Briefly, string phantoms were composed in agar mixed with intralipid (both Merck, UK) to mimic tissue-like scattering with red-coloured synthetic fibres (Smilco, USA) embedded at three different depths. PAI data were acquired at 100% laser energy with a 2kHz repetition rate. All animal procedures were conducted in accordance with project (PE12C2B96) and personal licenses (I544913B4, IA70F0365) issued under the United Kingdom Animals (Scientific Procedures) Act, 1986 and approved locally by the Cancer Research UK Cambridge Institute Animal Welfare Ethical Review Board.

To generate *in vivo* vascular tumour models, breast PDX tumour fragments were cryopreserved in a freezing media consisting of heat-inactivated foetal bovine serum (10500064, Gibco™, Fisher Scientific, Göteborg Sweden) and 10% dimethyl sulfoxide (D2650, Merck). The fragments were then defrosted at 37°C, washed with Dulbecco’s Modified Eagle Medium (41965039, Gibco™), mixed with matrigel (354262, Corning, NY, USA), and surgically implanted into the flank of 6–9 week-old NOD scid gamma (NSG) mice (#005557, Jax Stock, Charles River, UK), following standard protocols ^38,52^. The implantation involved one oestrogen receptor negative (ER-, n=6) PDX model and one oestrogen receptor positive (ER+, n=8) PDX model. After the tumours had reached an average diameter of ∼ 1 cm, the mice were imaged and then sacrificed, with the tumours collected in formalin for IHC analysis.

For the remaining breast cancer cell lines, seven-week old immuno-deficient female nude (BALB/c nu/nu) mice (Charles River) were inoculated orthotopically in the mammary fat pad of both flanks 1·100^6^ cells (either MCF7, n=7, or MDA-MB-231, n=6, random group assignment) in a final volume of 100 *μ*L of 1:1 phosphate-buffered saline (PBS, Gibco) and matrigel (BD). For MCF7, oestrogen implants (E2-M - 127 *β*-estradiol 90 days release, daily dose: 41.2-105.6 pg/ml, Belma Technologies) were implanted subcutanaously in the scruff of the neck 3 days before tumour cell injection.

For animal imaging, were mice anaesthetised using 3-5% isoflurane in 50% oxygen and 50% medical air. Mice were shaved and depilatory cream applied to remove hair that could generate image artefacts; single mice were placed into the PAI system, on a heat-pad maintained at 37°C. Respiratory rate was maintained between 70-80 bpm using isoflurane (∼ 1 − 2% concentration) throughout image acquisition. PAI data were acquired at 80% laser energy at 1kHz.

For string phantom imaging, phantoms were prepared following standard protocols ^74^ using agar mixed with intralipid (both Merck, UK) to mimic tissue-like scattering with red-coloured synthetic fibres (Smilco, USA) embedded at three different depths (top: 0.5 mm, middle: 1 mm, bottom: 2 mm).

### Immunohistochemistry

The tumour tissues, obtained for *ex vivo* validation, were processed by sectioning formalin-fixed paraffin-embedded (FFPE) samples. After deparaffinisation and rehydration, IHC analysis was performed on the tissues using the following antibodies: CD31 (anti-mouse 77699, Cell Signalling, London, UK), *α*SMA (anti-mouse ab5694, Abcam, Cambridge, UK), and carbonic anhydrase-IX (CAIX) (anti-human AB1001, Bioscience Slovakia, Bratislava, Slovakia), at concentrations of 1:100, 1:500, and 1:1000, respectively. The analysis was carried out using a BOND automated stainer, with a bond polymer refine detection kit (Leica Biosystems, Milton Keynes,UK), and 3,3’-diaminobenzadine as a substrate. The stained FFPE sections were scanned at a magnification of 20x using an Aperio ScanScope (Leica Biosystems, Milton Keynes, UK) and analysed with either ImageScope software or HALO Software (v2.2.1870, Indica Labs, Albuquerque, NM, USA). Regions of interest (ROIs) were drawn over the entire viable tumour area, and the built-in algorithms were customised to analyse the following: CD31 positive area (*μ*m^2^) normalised to the ROI area (*μ*m^2^) (reported as CD31 positivity (%)), area of CD31 positive pixels (*μ*m^2^) colocalised on adjacent serial section with *α*SMA positive pixels/CD31 positive area (*μ*m^2^).

### Datasets and Preprocessing

Image volume datasets were split into two categories: synthetic or experimental datasets (see Supplementary Table 6 for overview). The synthetic datasets were comprised of binary labels of 3D mathematically-generated vascular networks, which are paired with computer-simulated photoacoustic image volumes (n=449 for each). Here, physics-driven predictions are performed on the spatial architecture of the synthetic vasculature provided by the V-System in each binary image volume. The experimental datasets consisted of a string phantom (n=7), mouse ears (n=32), mouse skin (n=41) and breast cancer patient-derived xenograft (PDX) tumours in mice (n=445), and MCF7 and MDA-MB-231 tumours derived from bread cancer cell lines (n=204), all imaged *in vivo*. These datasets represent the *imaging domain* in VAN-GAN where *n* indicates the number of image volumes used. Note, the string phantoms were not used for training.

All photoacoustic datasets were stored as 32-bit greyscale 600 *×* 600 *×* 140 *μ*m^3^ (real) and 512 *×* 512 *×* 140 *μ*m^3^ (simulated) voxel tiff stacks with an isotropic voxel size of 20 *×* 20 *×* 20 *μ*m^3^ in the X-, Y- and Z-directions, where the Z-axis is perpendicular to the surface. All synthetic images were stored in an 8-bit format and generated with dimensions 512 *×* 512 *×* 140 *μ*m^3^ and an equal isotropic voxel size. As VAN-GAN trains on image subvolumes of size 128 *×* 128 *×* 128 *×* 1 voxels, all datasets were downsampled to 128 voxels in the Z-axis using a combination of maximum and bicubic downsampling, to ensure that depth-dependent SNR information is retained for training. All real and simulated photoacoustic images were normalised by performing XY slice-wise Z-score normalisation followed by thresholding of the top and bottom 0.05% of pixel intensities to correct for uneven illumination with respect to depth. Finally, datasets were normalised to a pixel intensity range of [−1, 1]. Datasets were partitioned with 10% assigned for testing and with the remaining 90% split 80/20 between training/validation.

### Network Training

Image patches were randomly sampled with a global batch size of two. Due to the sparsity of vessels with respect to the background for all datasets, a mapping function was applied when images were retrieved from the synthetic segmentation dataset. Here, the function detected whether a sampled volume contained any vessels to ensure VAN-GAN learnt how to segment vessels rather than just background. For 90% of sampled volumes, if no vessel was detected, a new image volume was sampled. In addition, to artificially extend the size of our datasets, all images were augmented via a rotation about the Z-axis randomly sampled from the set {0, *π/*2, *π*, 3*π/*2, 2*π*}.

All convolutional kernels were initialised using a He-normal initialiser and our loss functions hyperparameters were set to default values ^42,44^ *α* = 0.5, *λ* = 10 and *η* = *μ* = 5. Following Zhu et al. ^42^, training was performed for 200 epochs using the Adam optimizer ^75^ with a learning rate of 2 *×* 10^−4^ and 1^st^ and 2^nd^ moment estimates of 0.5 and 0.9, for both generators and discriminators. Each generator and discriminator was given a linear learning rate decay to 0 from the 100^th^ epoch.

Training was stabilised in two ways. Firstly, noise was applied to the real and fake input images to the discriminators ^62^. Noise was sampled from a Gaussian distribution with a mean and variance, *σ*, of 0.0 and 0.8^*i*^, where *i* was the i^th^ epoch, and so *σ* is annealed during training. Secondly, to avoid exploding gradients, a gradient clip normalisation strategy ^76^ where all gradients of each weight was individually clipped so that its norm is no higher than 100.

### Postprocessing

To reconstruct whole image volumes from generator output a sliding-window approach was employed ^34^. In summary, a sliding-window of size 128 *×* 128 *×* 128 voxels was strided across each inputted image with a stride length of 25 voxels in each XY direction. Output intensities were summed and the mean value calculated for each voxel location by tracking the number of times the window passed across a given voxel. To reduce edge artefacts, symmetric padding was used to ensure the sliding-window passed over each voxel for an equal number of instances. For segmenting an image, this reconstruction method results in a 32-bit 3D greyscale image where intensities indicate a probability that a voxel is a blood vessel. Consequently, following bicubic upsampling to an isotropic voxel size (140 voxels in the Z-axis), images were then thresholded based on the histogram of voxel intensities to binarise the image.

### Evaluation Metrics

Common machine learning metrics do not provide a complete picture of image segmentation performance for tubular-like structures ^77^. To evaluate our results, we calculated both standard segmentation metrics and a set of vascular descriptors ^38,78^ to provide a deeper insight into how well network morphology is predicted. The standard segmentation metrics used comprised of: F1 Score = 2 · (*precision* · *sensitivity*)*/*(*precision* + *sensitivity*); Intersection over Union (IoU) = *TP/*(*TP* + *FP* + *FN*); Sensitivity = *TP/*(*TP* + *FN*) and Specificity = *TN/*(*TN* + *FP*), where *TP* = true positive, *TN* = true negative, *FP* = false positive and *FN* = false negative.

To calculate vascular descriptors, all segmentations were skeletonised using the open-source package Russ-learn ^79,80^. The vascular skeletons allowed us to perform structural and topological data analyses on the vascular skeletons ^78,81^. The metrics were use are: number of vessels and branching nodes, vessel mean and standard deviation of diameters and lengths, network volume, surface density (the surface area of the vascular network normalised against the tissue volume), whereas topological descriptors consisted of connected components (or subnetworks, Betti-0) and looping structures (Betti-1) and network connectivity (the volume of the largest vascular subnetwork normalised against total network volume).

### Statistical Analysis

Statistical analyses were conducted using Prism (v9, GraphPad Software, San Diego, CA, USA). Comparisons of metrics between synthetic segmentations were computed using the non-parametric Friedman test. Comparisons of vascular descriptors for *in vivo* ear datasets were performed using either paired parametric or non-parametric t-tests depending on data satisfying normality. Comparison of vascular descriptors for *in vivo* tumour datasets were performed using Wilcoxon tests, with comparisons between ER- and ER+ or MCF7 and MDA-MB-231 tumour types made using unpaired non-parametric t-tests (Mann-Whitney tests). Statistical outliers were identified by five non-parametric tests: 1) Tukey’s fences; 2) Median Absolute Deviation (MAD); 3) Modified Z-Score; 4) percentiles (5^*th*^ and 95^*th*^ percentile cuttoffs) and 5) Hampel identifier. All P-values *<* 0.05 were considered statistically significant.

## Code Availability

All our software are open-source and available in Github repositories. VAN-GAN (2023, Version 1.0) [Computer software - https://github.com/psweens/VAN-GAN]. V-System (2022, Version 2.0) [Computer Software - https://github.com/psweens/V-System]. Vascular Topological Data Analysis (2022, Version 2.0) [Computer Software - https://github.com/psweens/Vascular-TDA].

## Data Availability

Scientific data supporting the findings of this study will be made available upon publication via the University of Cambridge Research Data Repository at: https://doi.org/10.17863/CAM.96379.

## Acknowledgements

PWS, TLL, LH, ELM, and SEB acknowledge the support from Cancer Research UK under grant numbers C14303/A17197, C9545/A29580, C47594/A16267, C197/A16465, C47594/A29448, and Cancer Research UK RadNet Cambridge under the grant number C17918/A28870. PWS acknowledges the support of the Wellcome Trust and University of Cambridge through an Interdisciplinary Fellowship under grant number 204845/Z/16/Z. TLL is supported by the Cambridge Trust. LH was funded from NPL’s MedAccel programme financed by the Department of Business, Energy and Industrial Strategy’s Industrial Strategy Challenge Fund. JG acknowledges funding from the Walter Benjamin Stipendium of the Deutsche Forschungsgemeinschaft. SEB is also funded by the EPSRC (EP/R003599/1). We thank the Cancer Research UK Cambridge Institute Biological Resources Unit and Imaging Core for their support in conducting this research. We also thank the laboratory of Prof. Carlos Caldas for their provision of the patient-derived xenograft material and support in establishing the models. For the purpose of open access, the corresponding author has applied a Creative Commons Attribution (CC BY) licence to any Author Accepted Manuscript version arising.

## Competing Interests

The authors have no conflict of interest related to the present manuscript to disclose.

## Author Contributions

Conceptualisation: PWS, SEB

Methodology: PWS, JG

Software: PWS

Validation: PWS, SEB

Formal Analysis: PWS

Investigation: PWS, LH

Resources: SEB

Data Collection: PWS, LH, TLL, ELB

Data Curation: PWS

Writing - original draft: PWS

Writing - review & editing: PWS, LH, TLL, ELB, JG, SEB

Visualisation: PWS

Project Administration: PWS, SEB

Funding Acquisition: PWS, SEB

## Dear Dr Pep Pàmies

We are pleased to submit our manuscript entitled ‘*Unsupervised segmentation of 3D microvascular photoacoustic images using deep generative learning’* for consideration for publication in *Nature Biomedical Engineering*. Our study introduces a novel multidisciplinary approach that leverages machine learning for the analysis of medical images obtained through photoacoustic imaging (PAI) to explore vascular structure and function. Our innovative solution stands to significantly benefit your readership by advancing research and diagnostics in fields that utilise PAI.

## Current Challenge

Microvascular network segmentation from image volumes presents a unique obstacle in biological research due to their complex 3D structures. This challenge is particularly evident in PAI, where photoacoustic artefacts significantly hinder the accurate and robust extraction of vascular parameters from high-resolution *in vivo* and *ex vivo* images, thus limiting their study in animal models and humans. While supervised deep neural networks have shown promise, the labour-intensive and error-prone nature of human annotations has limited their application to 3D images of blood vessels.

## Our Solution

To overcome this constraint, we introduce VAN-GAN (*Vessel Segmentation Generative Adversarial Network*), an unsupervised image-to-image translation framework, which eliminates the need for human-annotated ground-truth labels by leveraging mathematically derived synthetic blood vessel networks to segment 3D photoacoustic images.

> ***Our approach enables precise and unbiased blood vessel segmentations in 3D bioimages, eliminating the need for manual annotations***.

We demonstrate VAN-GAN’s superior performance in accurate and standardised segmentation of 3D vascular networks across a variety of *in silico, in vitro* and *in vivo* datasets, including human, patient-derived breast cancer xenograft (PDX) models.

## Key Novelties

- **Innovative approach for PAI segmentation**: VAN-GAN introduces a novel deep generative model tailored for 3D vascular network segmentation in PAI in the absence of human annotations.
- **Rivals supervised segmentation**: VAN-GAN challenges, for example, U-Net in segmenting 3D vasculatures from physics-derived photoacoustic image volumes.
- **Robustness to imaging artefacts**: VAN-GAN can account for PAI artefacts (e.g., depth-dependent SNR and illumination artefacts) which impede human annotators.
- **Versatility to features absent from training data**: ER+/ER-PDX and cell-line based models show that VAN-GAN can segment complex vasculatures that exceed the constraints imposed by its synthetic training data.

original research articles and reviews. We believe that the tuneability of VAN-GAN means that our approach is likely to have a broad impact in the study of vascular networks and biology using photoacoustics by providing a valuable tool to perform morphological analysis reliably and consistently across multi-modal bioimages. We are also convinced that the methodology of using physics-driven simulations in training means that the approach can readily extrapolate to other imaging modalities so will be of broad interest.

Given the novelty, significance, and broad interest of our findings to the fields of biomedical engineering, deep learning, and medical imaging, we believe that our manuscript is well-suited for publication in Nature Biomedical Engineering.

## Potential Reviewers

We would like to suggest the following expert reviewers for this manuscript:

1. Daniel Razansky, ETH/University of Zurich
  a.
  b. Expert in photoacoustics and prior research on deep learning in photoacoustic imaging.
2. Lena Maier-Hein, DKFZ
  a.
  b. Expert in data science and deep learning applied in photoacoustics.
3. Virginie Uhlmann, EMBL-EBI
  a.
  b. Expert in morphological bioimage analysis
4. Roger Zemp, University of Alberta
  a.
  b. Expert in clinical applications of novel biomedical imaging techniques.
5. Andreas Hauptmann, University of Oulu
  a.
  b. Expert in inverse problems in biomedical imaging.

I can confirm that the manuscript has been seen and approved by all authors. We have no conflict of interest to declare. This work has not been submitted for publication elsewhere and no results are reproduced from another source. We have had no prior discussions with any Nature Biomedical Engineering editor about the work descrived in our manuscript.

We appreciate your consideration of our study and look forward to hearing from you soon.

Yours sincerely,

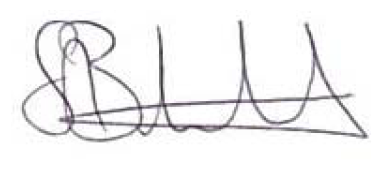

Prof. Sarah E Bohndiek, on behalf of all co-authors.

## Notes

### Competing Interest Statement

The authors have declared no competing interest.

### Summary of Updates

Manuscript adjusted to be more photoacoustic imaging focused with edits made throughout. Additions include ablation study and further comparisons to state-of-the-art segmentation methods.

