## Supplementary Information for "Unsupervised segmentation of 3D microvascular photoacoustic images using deep generative learning"

#### Supplementary Notes

##### Imaging Artefacts in Photoacoustic Mesoscopy

Photoacoustic mesoscopy is a non-invasive imaging technique that uses optical excitation and acoustic detection to produce high-resolution images of biological tissues. However, the technique is susceptible to imaging artefacts that can distort images and reduce image quality[1]. These artefacts arise from acoustic attenuation, acoustic reflection, optical scattering, acoustic diffraction, and thermal effects. Illumination artefacts, arise due to the limited top illumination of the field of view, leading to optical excitation and acoustic wave generation only in the upper part of absorbing structures that face the illumination and transducer array, resulting in inaccurate diameter estimations in the XZ plane compared to the XY plane[2]. Shadow artefacts, arise from obscuring objects, causing a signal loss in underlying objects due to strong optical or acoustic attenuation of the overlaying structure. Reflection artefacts, occur due to presence of acoustic reflectors/scatterers or strong acoustic reverberations of the absorbing object itself[3–5], leading to a signal echo near the object of interest. They can occur in-plane or out-of-plane depending on the position of the absorber/reflector[6].

#### A Comparison of Segmentation Methods for Tumour Vasculature

A comparison of performance between RF and VAN-GAN was performed for our PDX dataset (Supplementary Fig. 7A-C). Results were consist with prior analyses where VAN-GAN prediction significantly larger vascular density ($P < 0.0001$, Supplementary Fig. 7D) and vessel diameters ($P < 0.01$, Supplementary Fig. 7E). Vasculature segmented by VAN-GAN also exhibited greater connectivity compared to RF ($P < 0.001$, Supplementary Fig. 7F) resulting in a significant difference in computed vessel looping structures ($P < 0.001$, Supplementary Fig. 7G).

VAN-GAN allowed us to distinguish between structural differences between the ER+ and ER- subtypes, where ER+ tumours exhibited significantly larger mean blood vessel lengths ($P < 0.01$), which was not statistically significant using RF in Brown et al.[2] (Supplementary Fig. 8A). Similarly to Brown et al.[2], VAN-GAN predicts a significant difference in mean vessel diameters between the subtypes (Supplementary Fig. 8B), however, the trend is reversed with ER+ tumours on average exhibiting significantly larger diameters compared to ER- ($P < 0.01$). This highlights how VAN-GAN can provide an unbiased insight into biological characteristics of solid tumours.

### Supplementary Methods

#### V-System Stochastic Grammar

The aim of VAN-GAN is to learn the mappings between imaging and segmentation domains. Whilst datasets containing image volumes of real vascular networks are readily available, 3D blood vessel segmentations are not creating a need for realistic, 3D blood vessel segmentation labels for VAN-GAN to learn from. Mathematical angiogenesis models exist which incorporate biological mechanisms to simulate realistic, interconnected blood vessel networks[7], however, to generate a large library of vascular networks of a size applicable to deep learning can be computationally expensive.

Our synthetic vascular networks are generated using L-System stochastic definitions (termed here, *V-System*) that define a rule as *name* $\rightarrow$ *successor*, where symbols are replaced according to the respective rule[8,9]. Following Galarreta-Valverde et al.[9], rules are composed by symbols that have the following actions:

- $f(l, d)$: forward $l$ units in the direction vector and record vessel diameter, $d$;
- $+(\theta)$ and $-(\theta)$: rotate direction vector clockwise or anti-clockwise by $\theta^\circ$ on the plane

defined by the current direction vector;

- $/(\phi)$: rotate the current plane by $\phi^\circ$ about the direction vector;
- [: store the current state formed by the diameter;
- ]: restores the last saved state and allows the generation of a new branch;
- { and }: define the limits of a vessel segment.

For increasing number of recursive iterations, $n$, the string of instructions progressively becomes more complex. For example, in the case of modelling the growth of algae[8], using the grammatical rules A→ AB and A→ B, the axiom A becomes:

$n = 0$: A

$n = 1$: AB

$n = 2$: ABA

$n = 3$: ABAAB

$n = 4$: ABAABABA

The following string grammar was used to generate our synthetic vascular trees. Note, with the exception of $L(d_0)$ and the set of randomly sampled parameters, the following are not mathematical functions.

$$F \rightarrow S(n-1, d_0)[+(\theta_1)/(\phi_1)F(n-1, \theta_1)][-(\theta_2)/(\phi_2)F(n-1, d_2)], \tag{1}$$

where $d_1$ and $d_2$, and $\theta_1$ and $\theta_2$ are daughter branching diameters and bifurcation angles, respectively, stochastically calculated from previous studies[9–11]. The vessel tilt angles, $\phi_1$ and $\phi_2$, are sampled from a uniform distribution over the interval $[22.5°, 27.5°]$ multiplied by a random $\pm$ sign. Here, the string grammar $S$ is defined by:

$$S(n, d_0, \alpha, \beta) \rightarrow \begin{cases} \{S_0(n, d_0)\} & \text{if} \quad r < 0.5 \\ \{S_1(n, d_0)\} & \text{if} \quad r \geq 0.5 \end{cases}, \tag{2}$$

where $r$ is a random number generated over the interval $[0, 1)$. The function $S_i$ for $i = 0, 1$ is defined by

$$S_i(n, d_0) \rightarrow \begin{cases} D(n-1, \lambda_1 d_0) \pm (\theta_1) D(n-1, \lambda_2 d_0) & \\ \mp(\theta_2) D(n-1, \lambda_3 d_0) \mp (\theta_1) D(n-1, \delta_1 d_0) & \\ \pm(\theta_2) D(n-1, \delta_2 d_0) & \text{if} \quad r < 0.1 \\ D(n-1, d_0) \pm (\theta_1) D(n-1, d_0) \mp (\theta_2) D(n-1, d_0) & \\ \mp(\theta_1) D(n-1, d_0) \pm (\theta_2) D(n-1, d_0) & \text{if} \quad r \geq 0.1 \end{cases}, \qquad (3)$$

where $\lambda_j$ for $j = 1, 2, 3$ and $\delta_k$ for $k = 1, 2$ are growth and decay rates, respectively, sampled from the interval $[1, 1.5]$. We note that $\boldsymbol{\lambda}$ and $\boldsymbol{\delta}$ are vectors sorted in ascending and descending orders of magnitude, respectively. The string $S_i$ defines a branching vessel, which can randomly contain an aneurysm of random size (probability of 1 in 10).

$$D(n, d_0) \rightarrow f(L(d_0), d_0), \qquad (4)$$

where $L(d_0)$ the segment length function which randomly samples a value from the interval $[\epsilon d_0/5 - \Delta, \epsilon d_0/5 + \Delta]$. Here, the division by five is due to (3) where vessels are subdivided into five segments, and $\epsilon$ and $\Delta$ are randomly sampled from the intervals $[4, 10]$ and $[2, 4]$, respectively.

Using our grammatical rule, as an example, the axiom $F$ becomes:

$n = 0$: A

$n = 1$: {S}[+(38.3)/(0.0)F][-(36.7)/(0.0)F]

$n = 2$: {D+(0.0)D-(0.0)D-(0.0)D+(0.0)D}[+(37.4)/(25.5){S}[+(51.4)/(0.0)F][-(24.2)/(0.0)F]][-

(37.6)/(-25.7){S}[+(24.6)/(-24.0)F][-(51.0)/(-22.6)F]].

The resulting synthetic vascular network quantified for a set of statistical and topological vascular descriptors[2] in Supplementary Figure 6. Here, *V-System Recursion* defines the number of recursive iterations in which (1) is run. Exemplar vascular skeletons of increasing complexity are illustrated in Fig. 1 with distributions for our metrics given in Fig. 2.

#### Supervised Random Forest (RF) Pixel Classifier

Using the open source ilastik tool, users can pre-define a set of possible image features and manually annotate a subset of voxels in training images (either 2D or 3D images) for a random forest model to learn to recognise complex image features. The trained model can then be applied to the remaining unlabelled data and its accuracy improved through retrospective interactive training. Photoacoustic images were preprocessed for application to ilastik's random forest toolbox by applying a slice-wise median filter with a kernel size of $3 \times 3$ to remove noise and improve contrast. For each dataset, a sample of images were manually annotated with two label categories: blood vessel and tissue background. Here, approximately 10 XY images were labelled from each training image volume. All 3D colour / intensity, edge and texture features were selected for all Gaussian mean variational scales (0.3 to 10.0) to extract fine details at a small scale as well features from larger voxel neighbourhoods. RF models were trained by authors PWS, LH, TLL and ELB. For synthetic data, all vessel labels were provided for training along with a proportion of surrounding background labels (detailed in Brown et al.[2]). Trained models were applied to unlabelled images with predictions thresholded using default settings to create binary segmentation masks.

#### Model Training

VAN-GAN models were trained pragmatically as and when new datasets and computational hardware became available. Three VAN-GAN models were trained in total (see Supplementary Table 7). In each case, VAN-GAN predictions were thresholded at an intensity value of 0.5 (images scaled between 0 and 1.0) to create a segmentation mask. Increasing the batch size led to a noticeable improvement in VAN-GAN certainty between the vessel and tissue background. We observed that workstation (2), which employed a larger batch size of 4 (global batch size of 16), achieved better results compared to workstation (1) that used a batch size of 1 (global batch size of 2).

#### Comparing Vessel Diameters between Axial Planes

To compare the diameters of vessels in the axial $XY$- and $XZ$-planes, analysis was performed using MATLAB v2022b, MathWorks, Natick, MA, USA. Maximal intensity projections (MIPs) were created in each axial plane for each image. The diameters were analysed by creating a distance transform (using in-built function *bwdist*) to provide vessel diameters and vascular skeletons (using in-built function *bwmorph*). The resulting data provided a means to calculate mean vessel diameter for each plane. We note that MIPs were used as skeletonisation assumes radial symmetry, however, accuracy of this method declines for increasing vascular density of an image due to overlapping vessels when generating MIPs.

### Supplementary Figures

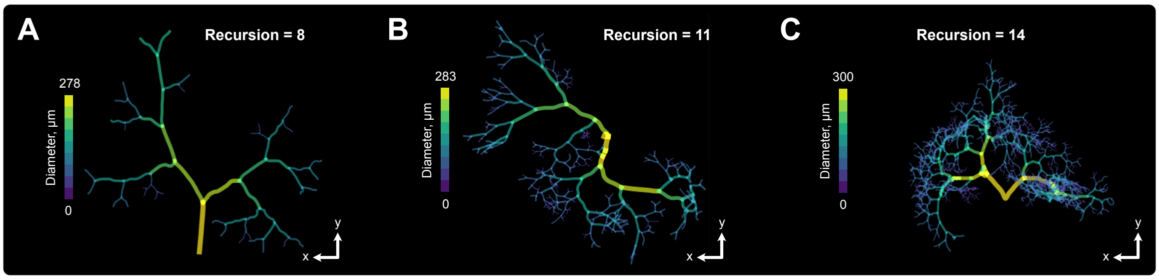


Figure 1: **Exemplar synthetic vasculatures of varying complexity.** 2D projections of networks, colour-coded to vessel diameters, are provided for increasing network complexity for (A) eight, (B) eleven and (C) fourteen grammatical recursions.

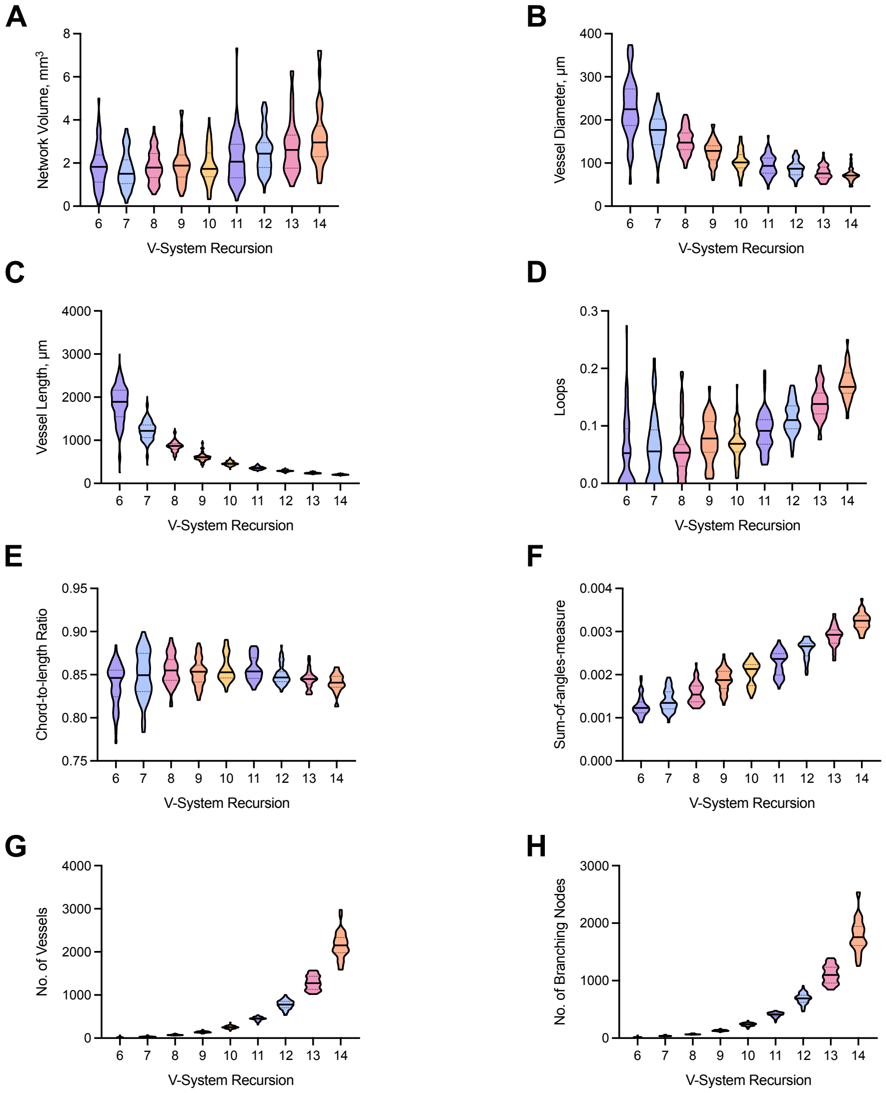


Figure 2: **Statistical and vascular descriptor distributions calculated from the ground truth synthetic vascular networks.** Truncated violin plots are separated into network complexity indicated by the number of V-System recursions. (A) Network volume, (B) mean vessel diameters, (C) mean vessel lengths, (D) mean number of network loops, (E) chord-to-length (CLR) ratio which defines vessel curvature, (F) sum-of-angles-measure (SOAM, a measure of vessel tortuosity), and the total number of (G) vessels and (H) branching nodes. Dashed and dotted lines indicate median, and upper (75$^{\text{th}}$) and lower (25$^{\text{th}}$) quartiles, respectively. Note, connected components and loops are normalised with respect to total number of edges per network.

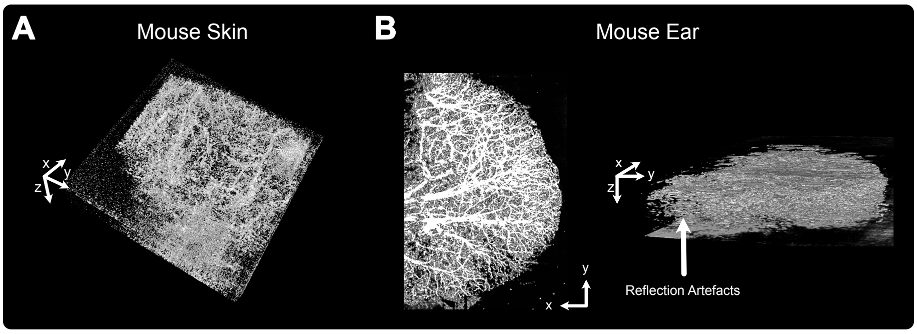


Figure 3: **Poor trained U-Net generalisability to *in vivo* data.** Application of our U-Net model trained on our simulated paired data to *in vivo* photoacoustic images of (A) mouse skin and (B) mouse ear. The U-Net model over segmented background noise such as reflection artefacts (shown in (B)).

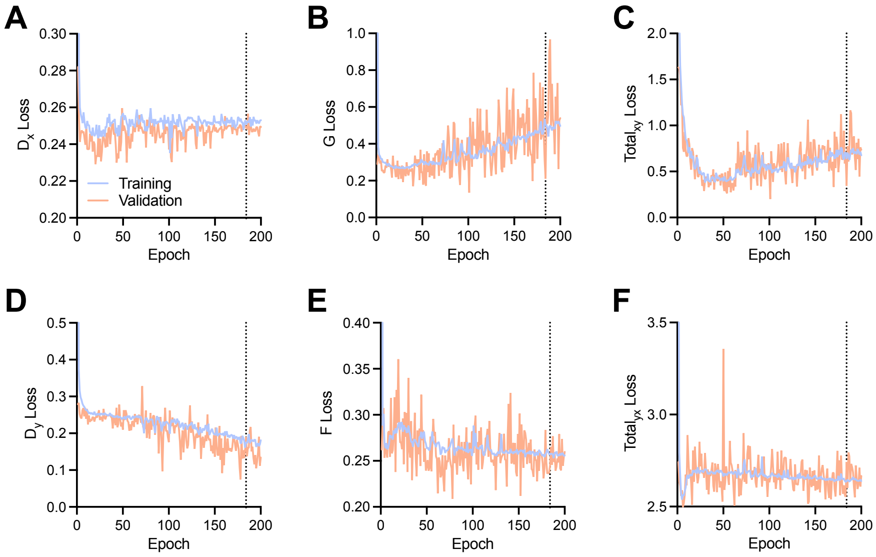


Figure 4: **VAN-GAN training and validation losses when applied to synthetic imaging and segmentation domains.** Training and validation losses are given per epoch for: discriminators (A) $D_x$ and (D) $D_y$; generators (B) $G$ and (E) $F$; and total losses (C) $x \rightarrow y$ and (F) $y \rightarrow x$. Dashed lines indicate the optimal epoch (epoch 184).

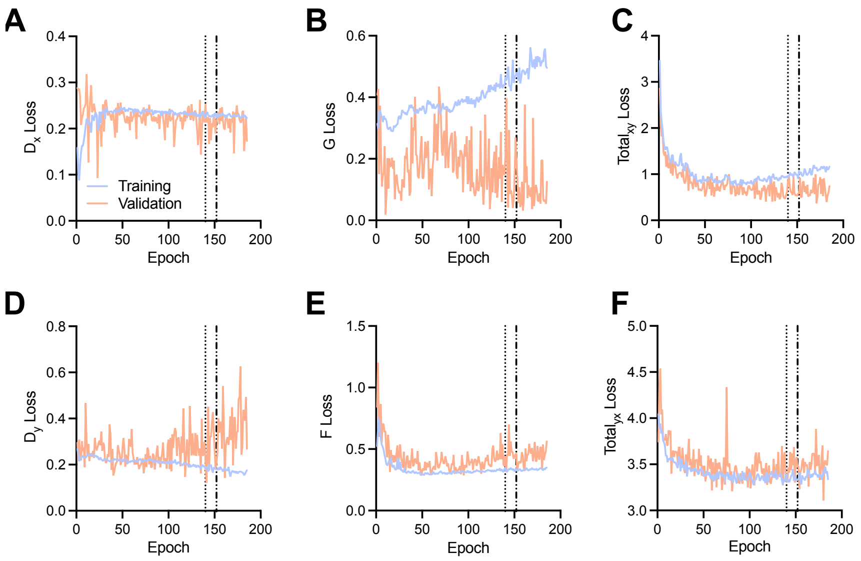


Figure 5: **VAN-GAN training and validation losses when applied to *in vivo* photoacoustic images of mouse skin, ear and breast cancer patient-derived xenograft tumours.** Training and validation losses are given per epoch for: discriminators (A) $D_x$ and (D) $D_y$; generators (B) $G$ and (E) $F$; and total losses (C) $x \rightarrow y$ and (F) $y \rightarrow x$. Dotted lines indicate epoch (140) used to segmented the ear dataset. Dashed lines indicate the optimal epoch (epoch 152). VAN-GAN experienced a vanishing gradient at the 185$^{\text{th}}$ epoch.

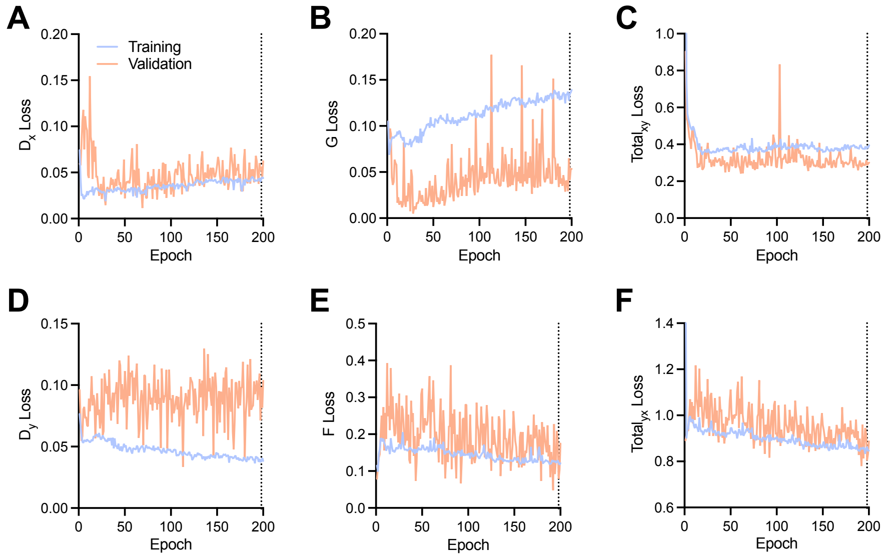


Figure 6: **VAN-GAN training and validation losses when applied to *in vivo* photoacoustic images of mouse skin, ear, breast cancer patient-derived xenograft tumours, and MCF7 and MDA-MB-231 tumours.** Training and validation losses are given per epoch for: discriminators (A) $D_x$ and (D) $D_y$; generators (B) $G$ and (E) $F$; and total losses (C) $x \rightarrow y$ and (F) $y \rightarrow x$. Dotted lines indicate epoch (140) used to segmented the ear dataset. Dashed lines indicate the optimal epoch (epoch 196).

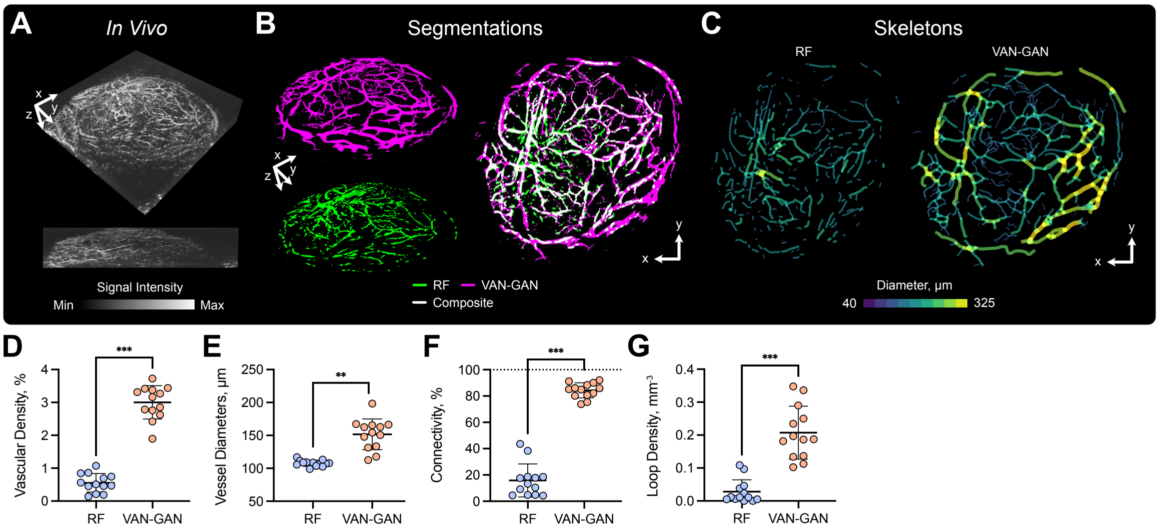


Figure 7: **VAN-GAN improves quantification of pathological vascular architecture.** Maximal intensity projections of: (A) *in vivo* signal intensity, (B) segmentation masks for (top left) VAN-GAN and (bottom left) the random forest pixel classifier (RF), and (right) a 2D overlay; and (C) vascular skeletons computed from (left) RF and (right) VAN-GAN segmentation predictions. We compute our vascular descriptors to compare results between the RF and VAN-GAN models. (D) Vascular density (the percentage of vascular volume with respect to tumour volume, %). (E) Mean vessel diameters ($\mu$m). (F) Connectivity (the volume percentage of the largest vascular subnetwork with respect to the sum of all subnetwork volumes, %). (G) Loop density (the number of network loops normalised against tumour volume, mm$^{-3}$). Mean and standard deviation of data are shown. Statistical significance indicated by * ($P < 0.05$), ** ($P < 0.01$), *** ($P < 0.001$) and **** ($P < 0.0001$).

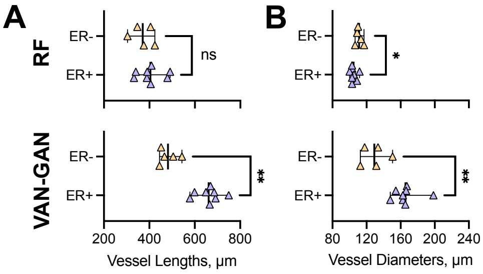


Figure 8: **Segmentation method can influence biological conclusions.** Computed mean vessel (A) lengths and (B) diameters for random forest classifier (RF, top) and VAN-GAN (bottom). Mean and standard deviation of data are shown. Statistical significance indicated by * ($P < 0.05$) and ** ($P < 0.01$) ($ns$ = not significant).

### Supplementary Tables

Table 1: **Network architecture for 3D deep residual U-Net.** Image input and output sizes are $128 \times 128 \times 128 \times 1$ (depth $\times$width $\times$height $\times$channel). *Conv i* represents the i$^{\text{th}}$ 3D convolutional layer. Note, for each encoding and decoding layer an identity mapping also exists in each residual block. Here, a 3D convolution of size $1 \times 1 \times 1$ is applied to the layer input and added to the layer output. The number of filters are equal to the other filters defined in each layer, and stride lengths are equal to 2 and 1 for encoding and decoding paths, respectively.

| | Level | Layer | Filters | Size | Stride | Output Size |
|---|---|---|---|---|---|---|
| Input | | | | | | $128 \times 128 \times 128 \times 1$ |
| Encoder | 1 | Conv 1 | 16 | $3 \times 3 \times 3$ | 1 | $128 \times 128 \times 128 \times 16$ |
| | | Conv 2 | 16 | $3 \times 3 \times 3$ | 1 | $128 \times 128 \times 128 \times 16$ |
| | 2 | Conv 3 | 32 | $3 \times 3 \times 3$ | 2 | $64 \times 64 \times 64 \times 32$ |
| | | Conv 4 | 32 | $3 \times 3 \times 3$ | 1 | $64 \times 64 \times 64 \times 32$ |
| | 3 | Conv 5 | 64 | $3 \times 3 \times 3$ | 2 | $32 \times 32 \times 32 \times 64$ |
| | | Conv 6 | 64 | $3 \times 3 \times 3$ | 1 | $32 \times 32 \times 32 \times 64$ |
| | 4 | Conv 7 | 128 | $3 \times 3 \times 3$ | 2 | $16 \times 16 \times 16 \times 128$ |
| | | Conv 8 | 128 | $3 \times 3 \times 3$ | 1 | $16 \times 16 \times 16 \times 128$ |
| Bridge | 5 | Conv 9 | 256 | $3 \times 3 \times 3$ | 1 | $8 \times 8 \times 8 \times 256$ |
| | | Conv 10 | 256 | $3 \times 3 \times 3$ | 1 | $8 \times 8 \times 8 \times 256$ |
| | | Conv 11 | 256 | $3 \times 3 \times 3$ | 1 | $8 \times 8 \times 8 \times 256$ |
| | | Conv 12 | 256 | $3 \times 3 \times 3$ | 1 | $8 \times 8 \times 8 \times 256$ |
| Decoder | 4 | Upsample 1 | - | $2 \times 2 \times 2$ | - | $16 \times 16 \times 16 \times 256$ |
| | | Conv 13 | 128 | $3 \times 3 \times 3$ | 1 | $16 \times 16 \times 16 \times 128$ |
| | | Conv 14 | 128 | $3 \times 3 \times 3$ | 1 | $16 \times 16 \times 16 \times 128$ |
| | 3 | Upsample 2 | - | $2 \times 2 \times 2$ | - | $32 \times 32 \times 32 \times 128$ |
| | | Conv 15 | 64 | $3 \times 3 \times 3$ | 1 | $32 \times 32 \times 32 \times 64$ |
| | | Conv 16 | 64 | $3 \times 3 \times 3$ | 1 | $32 \times 32 \times 32 \times 64$ |
| | 2 | Upsample 3 | - | $2 \times 2 \times 2$ | - | $64 \times 64 \times 64 \times 64$ |
| | | Conv 17 | 32 | $3 \times 3 \times 3$ | 1 | $64 \times 64 \times 64 \times 32$ |
| | | Conv 18 | 32 | $3 \times 3 \times 3$ | 1 | $64 \times 64 \times 64 \times 32$ |
| | 1 | Upsample 4 | - | $2 \times 2 \times 2$ | - | $128 \times 128 \times 128 \times 32$ |
| | | Conv 19 | 16 | $3 \times 3 \times 3$ | 1 | $128 \times 128 \times 128 \times 16$ |
| | | Conv 20 | 16 | $3 \times 3 \times 3$ | 1 | $128 \times 128 \times 128 \times 16$ |
| Output | | Conv 21 | 1 | $1 \times 1 \times 1$ | 1 | $128 \times 128 \times 128 \times 1$ |

Table 2: **Network architecture for 3D convolutional discriminator.** *Conv i* represents the $i^{th}$ 3D convolutional layer. Note, Gaussian noise was added prior to each convolution and spatial dropout subsequently applied post-convolution[12].

| | Conv Layer | Filters | Kernel Size | Stride | Dropout | Output Size |
|---|---|---|---|---|---|---|
| Input | | | | | | $128 \times 128 \times 128 \times 1$ |
| | Conv 1 | 64 | $4 \times 4 \times 4$ | 2 | 20% | $128 \times 128 \times 128 \times 64$ |
| | Conv 2 | 128 | $4 \times 4 \times 4$ | 2 | 20% | $64 \times 64 \times 64 \times 128$ |
| | Conv 3 | 256 | $4 \times 4 \times 4$ | 2 | 20% | $32 \times 32 \times 32 \times 256$ |
| | Conv 4 | 512 | $4 \times 4 \times 4$ | 2 | 20% | $16 \times 16 \times 16 \times 512$ |
| Output | Conv 5 | 1 | $3 \times 3 \times 3$ | 1 | 0% | $16 \times 16 \times 16 \times 1$ |

Table 3: **Network architecture for 3D U-Net.** Image input and output sizes are $128 \times 128 \times 128 \times 1$ (depth $\times$ width $\times$ height $\times$ channel). 3D convolutions and max pooling used.

| | Level | Layer | Filters | Kernel Size | Stride | Output Size |
|---|---|---|---|---|---|---|
| Input | | | | | | $128 \times 128 \times 128 \times 1$ |
| Encoder | 1 | Conv 1 | 32 | $3 \times 3 \times 3$ | 1 | $128 \times 128 \times 128 \times 16$ |
| | | Max Pooling 1 | - | $3 \times 3 \times 3$ | 2 | $128 \times 128 \times 128 \times 16$ |
| | 2 | Conv 3 | 64 | $3 \times 3 \times 3$ | 1 | $64 \times 64 \times 64 \times 32$ |
| | | Max Pooling 2 | - | $3 \times 3 \times 3$ | 2 | $64 \times 64 \times 64 \times 32$ |
| | 3 | Conv 5 | 128 | $3 \times 3 \times 3$ | 1 | $32 \times 32 \times 32 \times 64$ |
| | | Max Pooling 3 | - | $3 \times 3 \times 3$ | 2 | $32 \times 32 \times 32 \times 64$ |
| | 4 | Conv 7 | 256 | $3 \times 3 \times 3$ | 1 | $16 \times 16 \times 16 \times 128$ |
| | | Max Pooling 4 | - | $3 \times 3 \times 3$ | 2 | $16 \times 16 \times 16 \times 128$ |
| Bridge | 5 | Conv 9 | 512 | $3 \times 3 \times 3$ | 1 | $8 \times 8 \times 8 \times 256$ |
| | | Conv 10 | 512 | $3 \times 3 \times 3$ | 1 | $8 \times 8 \times 8 \times 256$ |
| Decoder | 4 | Upsample 1 | - | $2 \times 2 \times 2$ | - | $16 \times 16 \times 16 \times 128$ |
| | | Conv 11 | 256 | $3 \times 3 \times 3$ | 1 | $16 \times 16 \times 16 \times 128$ |
| | | Conv 12 | 256 | $3 \times 3 \times 3$ | 1 | $16 \times 16 \times 16 \times 128$ |
| | 3 | Upsample 2 | - | $2 \times 2 \times 2$ | - | $32 \times 32 \times 32 \times 128$ |
| | | Conv 13 | 128 | $3 \times 3 \times 3$ | 1 | $32 \times 32 \times 32 \times 64$ |
| | | Conv 14 | 128 | $3 \times 3 \times 3$ | 1 | $32 \times 32 \times 32 \times 64$ |
| | 2 | Upsample 3 | - | $2 \times 2 \times 2$ | - | $64 \times 64 \times 64 \times 64$ |
| | | Conv 15 | 64 | $3 \times 3 \times 3$ | 1 | $64 \times 64 \times 64 \times 32$ |
| | | Conv 16 | 64 | $3 \times 3 \times 3$ | 1 | $64 \times 64 \times 64 \times 32$ |
| | 1 | Upsample 4 | - | $2 \times 2 \times 2$ | - | $128 \times 128 \times 128 \times 32$ |
| | | Conv 17 | 32 | $3 \times 3 \times 3$ | 1 | $128 \times 128 \times 128 \times 16$ |
| | | Conv 18 | 32 | $3 \times 3 \times 3$ | 1 | $128 \times 128 \times 128 \times 16$ |
| Output | | Conv 19 | 1 | $1 \times 1 \times 1$ | 1 | $128 \times 128 \times 128 \times 1$ |

Table 4: **CycleGAN generator architecture.** Each residual block (RB) is of the same type as used in VAN-GAN, consisting of of two 3D convolutional layers with a kernel size of $3 \times 3 \times 3$ and stride of $1 \times 1 \times 1$.

| | Level | Layer | Filters | Kernel Size | Stride | Output Size |
|---|---|---|---|---|---|---|
| Input | | | | | | $128 \times 128 \times 128 \times 1$ |
| Downsampling | 1 | Conv 1 | 32 | $7 \times 7 \times 7$ | 1 | $128 \times 128 \times 128 \times 32$ |
| | 2 | Conv 2 | 64 | $3 \times 3 \times 3$ | 2 | $64 \times 64 \times 64 \times 64$ |
| | 3 | Conv 3 | 128 | $3 \times 3 \times 3$ | 2 | $32 \times 32 \times 32 \times 128$ |
| | 4 | Conv 4 | 256 | $3 \times 3 \times 3$ | 2 | $16 \times 16 \times 16 \times 256$ |
| Residual Blocks | 5 | RB 1 | 256 | $3 \times 3 \times 3$ | 1 | $16 \times 16 \times 16 \times 256$ |
| | 6 | RB 2 | 256 | $3 \times 3 \times 3$ | 1 | $16 \times 16 \times 16 \times 256$ |
| | 7 | RB 3 | 256 | $3 \times 3 \times 3$ | 1 | $16 \times 16 \times 16 \times 256$ |
| | 8 | RB 4 | 256 | $3 \times 3 \times 3$ | 1 | $16 \times 16 \times 16 \times 256$ |
| | 9 | RB 5 | 256 | $3 \times 3 \times 3$ | 1 | $16 \times 16 \times 16 \times 256$ |
| | 10 | RB 6 | 256 | $3 \times 3 \times 3$ | 1 | $16 \times 16 \times 16 \times 256$ |
| Upsampling | 11 | Upsample 1 | 128 | $2 \times 2 \times 2$ | 1 | $32 \times 32 \times 32 \times 256$ |
| | | Conv 5 | 32 | $4 \times 4 \times 4$ | 1 | $32 \times 32 \times 32 \times 128$ |
| | 12 | Upsample 2 | 64 | $2 \times 2 \times 2$ | 1 | $64 \times 64 \times 64 \times 128$ |
| | | Conv 6 | 32 | $4 \times 4 \times 4$ | 1 | $64 \times 64 \times 64 \times 64$ |
| | 13 | Upsample 3 | 32 | $2 \times 2 \times 2$ | 1 | $128 \times 128 \times 128 \times 64$ |
| | | Conv 7 | 32 | $4 \times 4 \times 4$ | 1 | $128 \times 128 \times 128 \times 32$ |
| Output | | Conv 8 | 1 | $7 \times 7 \times 7$ | 1 | $128 \times 128 \times 128 \times 1$ |

Table 5: **Evaluating segmentation performance against 2D maximal intensity projections.** Segmentation masks are calculated for a 3D image using two random forest pixel classifiers (RF) trained separately by two experts (indicated by subscripts *1* and *2*) and our VAN-GAN method. Maximal intensity projections were calculate for each segmentation prediction for a direct comparison against the 2D ground truth masks using F1 Score, intersection over union (IoU), sensitivity and specificity performance metrics. Note, two scores per metric indicates that the metric was computed against the the ground truth generated by user (left) one or (right) two. The highest metric score per ground truth is shown in bold.

| Dataset | Method | F1 Score | IoU | Sensitivity | Specificity |
|---|---|---|---|---|---|
| **Mouse Ear** | $RF_1$ | **0.658/0.668** | **0.491/0.501** | 0.721/0.694 | 0.905/**0.922** |
| | $RF_2$ | 0.633/0.623 | 0.463/0.455 | **0.785/0.729** | 0.860/0.869 |
| | VAN-GAN | 0.649/0.630 | 0.480/0.460 | 0.688/0.636 | **0.912/0.922** |
| **Mouse Skin** | $RF_1$ | **0.584/0.613** | **0.413/0.443** | 0.755/0.711 | 0.858/0.879 |
| | $RF_2$ | 0.562/0.604 | 0.391/0.434 | **0.769/0.739** | 0.831/0.855 |
| | VAN-GAN | 0.529/0.533 | 0.360/0.364 | 0.580/0.532 | **0.897/0.911** |

Table 6: Mesoscopic photoacoustic image dataset details. Note, voxel size and image dimensions are defined by $X \times Y \times Z$ where the Z-axis is perpendicular to the skin surface. Dataset size indicates the number of image volumes. PA = photoacoustic and PDX = breast cancer patient-derived xenograft.

| Dataset | Voxel Size ($\mu m^3$) | Image Dimensions | Dataset Size |
|---|---|---|---|
| **Synthetic** | | | |
| Synthetic Vasculature | $20 \times 20 \times 20$ | $512 \times 512 \times 140$ | 449 |
| PA Simulations | $20 \times 20 \times 20$ | $512 \times 512 \times 140$ | 449 |
| **Experimental** | | | |
| Mouse Ear | $20 \times 20 \times 20$ | $600 \times 600 \times 140$ | 32 |
| Mouse Skin | $20 \times 20 \times 20$ | $600 \times 600 \times 140$ | 41 |
| PDX Tumours | $20 \times 20 \times 20$ | $600 \times 600 \times 140$ | 445 |
| MCF7 & MDA-MB-231 Tumours | $20 \times 20 \times 20$ | $600 \times 600 \times 140$ | 204 |

Table 7: Trained VAN-GAN models with corresponding datasets and workstation specifications used. Workstations are labelled by: 1) a Dell Precision 7920T with a Dual Intel Xeon Gold 5120 CPU with 128GB RAM and two NVIDIA Quadro GV100 32GB GPUs with NVLink; 2) a custom built workstation with a Intel Xeon Gold 5220 CPU with 128GB RAM and four NVIDIA RTX A6000 48GB GPUs with NVLinks. Note, the synthetic vascular dataset is used in all models. PA = photoacoustic and PDX = breast cancer patient-derived xenograft tumours.

| VAN-GAN Model | Datasets Used | Workstation | Training Time | Figures |
|---|---|---|---|---|
| 1 | PA Simulations | 1 | 54 hours | Fig. 2, 3, 5B,C,F,G |
| 2 | Mouse Ear<br>Mouse Skin<br>PDX | 1 | 76 hours | Fig. 4, 5B,C-E,G, 6A-D |
| 3 | Mouse ear<br>Mouse Skin<br>PDX<br>MCF7 & MDA-MB-231 | 2 | 36 hours | Fig. 6E-I |
